# Early life exposure to cigarette smoke primes lung function and DNA methylation changes at *Cyp1a1* upon exposure later in life

**DOI:** 10.1101/2023.03.03.530858

**Authors:** Chinonye Doris Onuzulu, Samantha Lee, Sujata Basu, Jeannette Comte, Yan Hai, Nikho Hizon, Shivam Chadha, Maria Shenna Fauni, Shana Kahnamoui, Bo Xiang, Andrew J. Halayko, Vernon W. Dolinsky, Christopher Pascoe, Meaghan J. Jones

## Abstract

Prenatal and early life exposure to cigarette smoke (CS) have repeatedly been shown to induce stable, long-term changes in DNA methylation (DNAm) in offspring. It has been hypothesized that these changes might be functionally related to the known outcomes of prenatal and early life CS exposure, which include impaired lung development, altered lung function and increased risk of asthma and wheeze. However, to date, few studies have examined DNAm changes induced by prenatal CS in tissues of the lung, and even fewer have attempted to examine the specific influences of prenatal versus early postnatal exposures. Here, we have established a mouse model of CS exposure which isolates the effects of prenatal and early postnatal CS exposures in early life. We have used this model to measure the effects of prenatal and/or postnatal CS exposures on lung function and immune cell infiltration as well as DNAm and expression of *Cyp1a1*, a candidate gene previously observed to demonstrate DNAm differences upon CS exposure in humans. Our study revealed that exposure to CS prenatally and in the early postnatal period cause long-lasting differences in offspring lung function, gene expression and lung *Cyp1a1* DNAm, which wane over time but are reestablished upon re-exposure to CS in adulthood. This study creates a testable mouse model which can be used to investigate the effects of prenatal and early postnatal CS exposures and will contribute to the design of intervention strategies to mediate these detrimental effects.

## Introduction

The developmental origins of health and disease (DOHaD) hypothesis postulates that environmental insults in the intrauterine and early postnatal environment program the developing fetus through changes in cellular function and structure, and that these changes can last through adulthood^1, 2^. Numerous studies have shown that factors such as maternal nutrition, infections, stress and environmental exposure to toxins such as cigarette smoke (CS) can in fact produce long-lasting health consequences in offspring^3, 4^. CS is a common environmental toxin in humans with about 9.5% of women of reproductive age (18-34 years) reported as active tobacco smokers in Canada in 2021^5^. Cigarette smoke is made up of thousands of components which have detrimental effects, some of which can also cross the placenta and affect offspring development^6–10^.

Early life CS exposure alters offspring growth and development, lung function, and is associated with airway hyperresponsiveness, asthma and wheeze in children^11–13^. In some cases, these outcomes persist into adulthood, suggesting that early life CS ‘leaves a mark’ on the offspring, a process that has been called *biological embedding*^14–16^. Epigenetic modifications have been identified as potential mechanisms by which biological embedding takes place^16, 17^. The three main components of the epigenome – DNA methylation (DNAm), histone modifications, and non-coding RNAs^18^ - are dynamic and reflect past environmental exposures, meaning that environmental insults in early life can elicit changes in epigenetic marks which cells would propagate over time, and thus could translate to disease phenotypes in later life.

CS exposure *in utero* has been associated with differential DNAm across thousands of genes in exposed children, suggesting that DNAm changes might directly or indirectly link early life CS exposure with later life outcomes^19–21^ ^21–25^. While most epigenome-wide association studies (EWAS) investigating association between maternal smoking and DNAm have been conducted in newborn umbilical cord blood^19–25^, others have also reported differential DNAm in placenta^26^, nasal epithelia^27^, buccal cells and peripheral blood of adolescents or adults^28–30^. The largest EWAS to date which investigated effects of prenatal smoke exposure on umbilical cord blood DNAm was a meta-analysis across 13 cohorts (n = 6685) in the Pregnancy and Childhood Epigenetics Consortium (PACE)^20^. Results from the PACE study reported differential methylation at over 6000 CpGs, with some of the top sites being Aryl Hydrocarbon Receptor Repressor (*AHRR*), Cytochrome P450 1A1 (*CYP1A1*), Myosin 1G (*MYO1G*) and Growth Factor Independent 1 Transcriptional Repressor (*GFI1*). These genes have also been associated with maternal smoking in other similar EWAS studies^19, 20, 23, 31^. They are important in detoxification, apoptosis and cell proliferation, and have been linked to the development of orofacial clefts, oncogenesis and asthma^19, 20, 31^. Existing studies including PACE have not been able to investigate the effects of prenatal smoking after birth, nor could they measure DNAm in tissues that might be more proximal to CS-mediated disease, such as the lungs.

While there have been human studies outlining the long-term effects of prenatal CS exposure on offspring, one challenge of these studies is their inability to effectively account for the effects of the postnatal environment. One study which measured the long-term effects of prenatal CS on offspring lung function found no significant association between second-hand smoke exposure while pregnant and long-term offspring lung function after controlling for postnatal smoke exposure^32^. It is difficult in human studies to completely rule out the effect of postnatal smoke exposure since prenatal and postnatal exposures are typically very similar^33, 34^. Therefore, there is a need to investigate the differences between DNAm signatures and health outcomes observed due to prenatal versus postnatal CS exposure, especially since they have been shown to produce mixed/confounding effects^35–38^. Animal models provide a solution to these problems by allowing the study of the effects of prenatal versus postnatal CS exposure, and enable research into longitudinal, cross-tissue and sex-specific effects of early life CS exposure.

Here, we have created a mouse model to study the effects of prenatal, postnatal, and combined early life CS on offspring DNAm and lung phenotype. Our model successfully recapitulates some of the phenotypes identified in humans exposed to prenatal CS and provides further information on the dynamics of early life CS exposure. We show that early life CS exposure alters offspring DNAm at *Cyp1a1*, lung function, and gene expression in patterns specific to their early life exposure periods. Critically, we show that the early life period can create lasting molecular memories, and that exposure to CS in adulthood recapitulates patterns set by early life CS exposure.

## Materials and Methods

### Animals

This experiment was approved by the Animal Research Ethics and Compliance Committee of the University of Manitoba. Balb/C mice (Charles River Laboratories, Massachusetts, United States) were supplied with standard laboratory chow and clean water *ad libitum* except during exposures and housed, four mice of a single sex per cage unless otherwise noted, in individually ventilated cages in a 12-hour light/dark cycle throughout the experiment.

### Development of murine model of early life CS exposure

Standard 1R6F research cigarettes (University of Kentucky, Lexington, KY) were used for this experiment. CS was delivered via the SCIREQ InExpose smoking robot (SCIREQ, Montreal, QC, Canada). Female mice were left to acclimate for 1 week, then divided into 2 groups: 16 control and 16 CS-exposed mice. Beginning at 8 weeks, control mice were exposed to room air in a foreign cage, and CS mice exposed to whole body CS twice daily. Total estimated particulate matter per certificate of analysis from the cigarette manufacturer was 46.8 mg/cigarette, administered with a flow rate of 2L/min and one puff per minute, with 6 puffs per cigarette. CS mice were exposed for 9 weeks, beginning with a pre-pregnancy period of 3 weeks, after which they were mated (2 females to 1 male) with male mice. Once a vaginal plug was achieved, we separated female mice into individual cages for the rest of the experiment. We continued all smoke and room air exposures on CS and control mice respectively, throughout mating, pregnancy, birth, and stopped at weaning (3 weeks post-birth). At birth, we culled offspring to 6 offspring per litter, and cross-fostered half of CS offspring with half of control offspring, matching offspring by birth date to reduce potential of rejection. Pups were marked with toe tattoos to identify birth group. Using our cross-fostering strategy, we generated 4 distinct offspring groups: control with no CS exposure (“Con”), offspring exposed to prenatal CS only (“Pre”), offspring exposed to postnatal CS only (“Post”), and offspring that received both prenatal and postnatal CS (“Full”). Throughout the experiment, offspring were never directly exposed to CS, but were indirectly exposed *in utero*, or via breastmilk or fur from dams. We sacrificed and collected tissues from offspring not cross-fostered at birth. Dams and offspring were weighed weekly. Three weeks after birth, offspring were weaned, CS exposures stopped, and dams underwent lung function testing followed by tissue collection. Pups were housed by sex, 4-6 to a cage. While original plans were to assess phenotype at 8 weeks of age, due to SARS-CoV2-related lab lockdowns animals were maintained until 16-20 weeks of age, after which lung function and tissue collection were conducted.

### CS re-exposure experiment

Female offspring which were not euthanized at 16-20 weeks were left undisturbed until 60 weeks of age. Half of them were re-exposed for 2 hours to full body CS for 5 consecutive days per week, for 21 days in total. The other half which was not re-exposed to CS served as the adult control group. Lung function, lavage and tissue collection were performed as described.

### Lung function measurement

As previously described^39, 40^, mice were anesthetized with sodium pentobarbital and lung function measured using the SCREQ Flexivent small animal ventilator (SCIREQ Inc., Montreal, QC, Canada). Total airway resistance (Rrs), Newtonian resistance (Rn), tissue resistance (G) and elastance (H) were measured at baseline in response to nebulized saline, and changes in these parameters were also measured in response to increasing concentrations of nebulized methacholine (3 to 50 mg/ml). Using a three-parameter logistic regression, we fitted a dose-response curve and then calculated the slope of the curve in order to assess overall methacholine sensitivity^41^ in the lungs after CS exposure.

### Bronchoalveolar lavage

Following lung function measurement, mouse lungs were washed twice through a tracheal cannula with 1 ml of phosphate buffer saline (PBS) each wash. Bronchoalveolar lavage fluid (BALF) was then centrifuged at 4°C at 1200 rpm for 10 minutes to obtain cell-free supernatants which were stored at - 20°C for later cytokine analysis. Cell pellets were resuspended in 1 ml of PBS and total cell counts were performed using a hemocytometer. Differential cell counts were performed by placing 100ul of resuspended pellets on glass slides using cytospin columns, and staining cells using a modified Wright-Giemsa stain (Hema 3 Stat Pack). Cell differentials were counted using a Carl Zeiss Axio Observer ZI microscope.

### Blood and tissue collection and preparation

Whole blood was collected from dams at pup weaning and from offspring at 16 and 63 weeks. Blood was collected by severing the abdominal aorta, into EDTA-coated tubes, centrifuged at 4000 rpm for 15 minutes to obtain plasma. Blood cell pellets and plasma were snap-frozen in dry ice and stored at - 80°C for future measurements. Left, inferior, postcaval and middle and superior lung lobes were collected separately and snap-frozen in dry ice for DNAm and gene expression analyses.

### DNA/RNA isolation from lungs and blood

DNA and RNA were extracted from mouse tissues using the Invitrogen DNA and RNA isolation kits respectively. Left lungs were homogenized using the Qiagen Tissue Lyser II, followed by simultaneous DNA and RNA isolation using same Invitrogen DNA and RNA isolation kits. DNA and RNA were quantified using a NanoDrop spectrophotometer (NanoDrop Technologies, USA).

### Selection of mouse candidate genes to measure DNAm

To obtain candidate genes at which to measure DNAm, we selected 2 of the top differentially methylated CpGs from the meta-analysis conducted in human cohorts^20^. The 2 chosen human CpGs with some of the largest effect sizes were selected from genome build GRCh37/hg19: *AHRR* at position chr5:373378 (cg05575921) and *CYP1A1* at position chr15:75019251 (cg22549041). We then blasted the resulting human sequence against the *Mus musculus* genome assembly on NCBI. Using R studio version 3.6, we performed a muscle^42^ alignment between human and resulting mouse sequences to detect mouse CpGs which exactly aligned with or were closest in position to the human CpGs. The 2 mouse CpGs were selected from genome build GRCm38/mm10 based on results of the alignment: position chr13:74260517 for *Ahrr* and position chr9:57696335 for *Cyp1a1.* Importantly, the *Ahrr* gene is not conserved between human and mice, and while we selected the closest mouse CpG, it may not be comparable to the human position. To select a CpG position for the mouse control gene, we selected a human CpG position which was not differentially methylated in newborn cord blood (human genome GRCh38/hg38, *PRKAA1* cg13345558, position chr5:40796738), as reported in the PACE study. We then identified a corresponding mouse CpG (genome build GRCm39/mm39 position chr15:5174566) as described above.

### Measurement of candidate gene DNAm

500 ng of DNA isolated from the left lungs was treated with sodium bisulfite (Zymo Research) to generate bisulfite-converted DNA (bcDNA), following the manufacturer’s protocol. Following conversion, bcDNA was then amplified by PCR using the following conditions: 95°C for 15 min, followed by 45 cycles of 95°C for 30 secs, 58°C for 30 secs, 72°C for 30 secs, and then followed by 72°C for 5 min. Mouse *Ahrr* and *Cyp1a1* primers were generated using the PyroMark Assay Design software version 2.0 (Qiagen). Primer sequences can be found in Table S1. DNAm at candidate genes was measured using the Qiagen Q48 pyrosequencer. Prior to DNAm measurement in samples, CpG assays were validated in duplicates using mixtures of completely methylated and completely unmethylated control DNA (0%, 25%, 50%, 75%, and 100% methylated). After validation, sample DNAm was then measured using the optimized assay conditions.

### Gene expression measurement

200ng of lung mRNA was treated with ezDNase (Thermo Fisher Scientific, Inc.) and then converted to cDNA using Maxima cDNA synthesis Kit (Thermo Fisher Scientific, Inc.) according to the manufacturer’s protocol. Quantitative real-time RT-PCR (qPCR) was performed on the QuantStudio 3 Real-Time PCR System (Thermo Fisher Scientific, Waltham, MA), using the SYBR-Green Master Mix (Applied Biosystems; Thermo Fisher Scientific, Inc.) to determine the relative expression levels of *Ahrr* and *Cyp1a1* in lung tissues. β*-actin* and *Eif2a* were used as reference and normalization controls. Samples were run in duplicates under the following cycling conditions: Holding stage, 1 cycle of 50°C for 2 min and 95°C for 2 min; cycling stage, 40 cycles of 95°C for 15 sec, 57°C for 15 sec and 72°C for 1 min; and melt curve stage, 95°C for 15 sec, 60°C for 1 min and 95°C for 15. The primer sequences used can be found in Table S2. *Ahrr* and *Cyp1a1* mRNA levels were quantified using the ΔΔ^Cq^ method^43, 44^ and normalized against mean β*-actin* and *Eif2a* levels in the same sample.

### Measurement of the effects of cross-fostering

Since we cross-fostered offspring at birth to create our model for early life CS exposure, we needed to rule out any potential effects of the cross-fostering process on offspring. 4 female and 2 male Balb/C mice were purchased at 6 weeks of age, separate from the mice used for the major experiment. They were left to acclimate for 2 weeks, fed and housed as described above. At 8 weeks of age, mice were placed together (2 females to 1 male) and mated for 3 days. At birth, we culled offspring to 6 offspring per litter, and cross-fostered half of resulting offspring, matching offspring by birth date to reduce potential of rejection. Pups were marked with a toe tattoo to identify birth group. Cross-fostering led to the generation of 2 groups of offspring: non cross-fostered and cross-fostered groups. We performed lung lavage, sacrificed and collected tissues from all offspring at 8 weeks of age.

### Statistical analyses

Statistical analysis was done using R (version 3.6.1). Percent DNAm at each CpG was averaged across duplicates and between-group differences were analyzed using a one-way ANOVA. Lung function and differential cell count data were analyzed using two-way ANOVA, and multiple comparison was performed between groups at each methacholine dose. *P* values < 0.05 were considered significant.

Adjustment for litter size effects was conducted on significant values using simple linear regression. Figures were also produced in R, using the *ggplot2* package.

## Results

### Development of a protocol to study the effects of early life CS exposures

We created a mouse model to study the effects of early life CS exposure on offspring, and specifically, to isolate prenatal and postnatal CS exposures. Exposure of Balb/C dams to whole-body CS was carried out for 9 weeks, beginning 3 weeks before mating and pregnancy, and lasting until 3 weeks after birth (Figure 1A). To isolate the effects of prenatal and postnatal-only smoke exposures, we cross-fostered half of the offspring born to control and smoke-exposed dams at birth, with toe tattoos to identify cross fostered offspring (Figure 1A). All cross-fostering was carried out no more than 24 hours after birth of matched control and CS-exposed litters. Only one litter, including both cross fostered and not cross fostered offspring, was lost. All smoke-exposed dams tolerated smoke exposures well, and our exposure paradigm did not result in loss of any dams. Since mating resulted in variable plug/conception dates, we ensured that all analyses reported here were carried out on offspring born no more than 48 hours apart.

Maternal smoking can reduce litter size^45, 46^ and pup birth weight in mice, which might have indirect effects on pup development^46, 47^. To ensure that our DNAm and phenotypic analyses are not confounded by such effects, we tested the effects of our smoking model on offspring birth weight and litter size. We found a slightly larger litter size in control dams (Mean = 6.56, SD = 1.46) than smoke-exposed ones (Mean = 5.38, SD = 1.36) (Figure 1B, T-test *p* = 0.024). Birthweight in male and female animals was slightly but not significantly larger in CS than control pups, perhaps due to the smaller litter size (Figure 1C). We performed sensitivity tests in downstream analyses to identify the potential effects of litter size.

**Figure 1:**
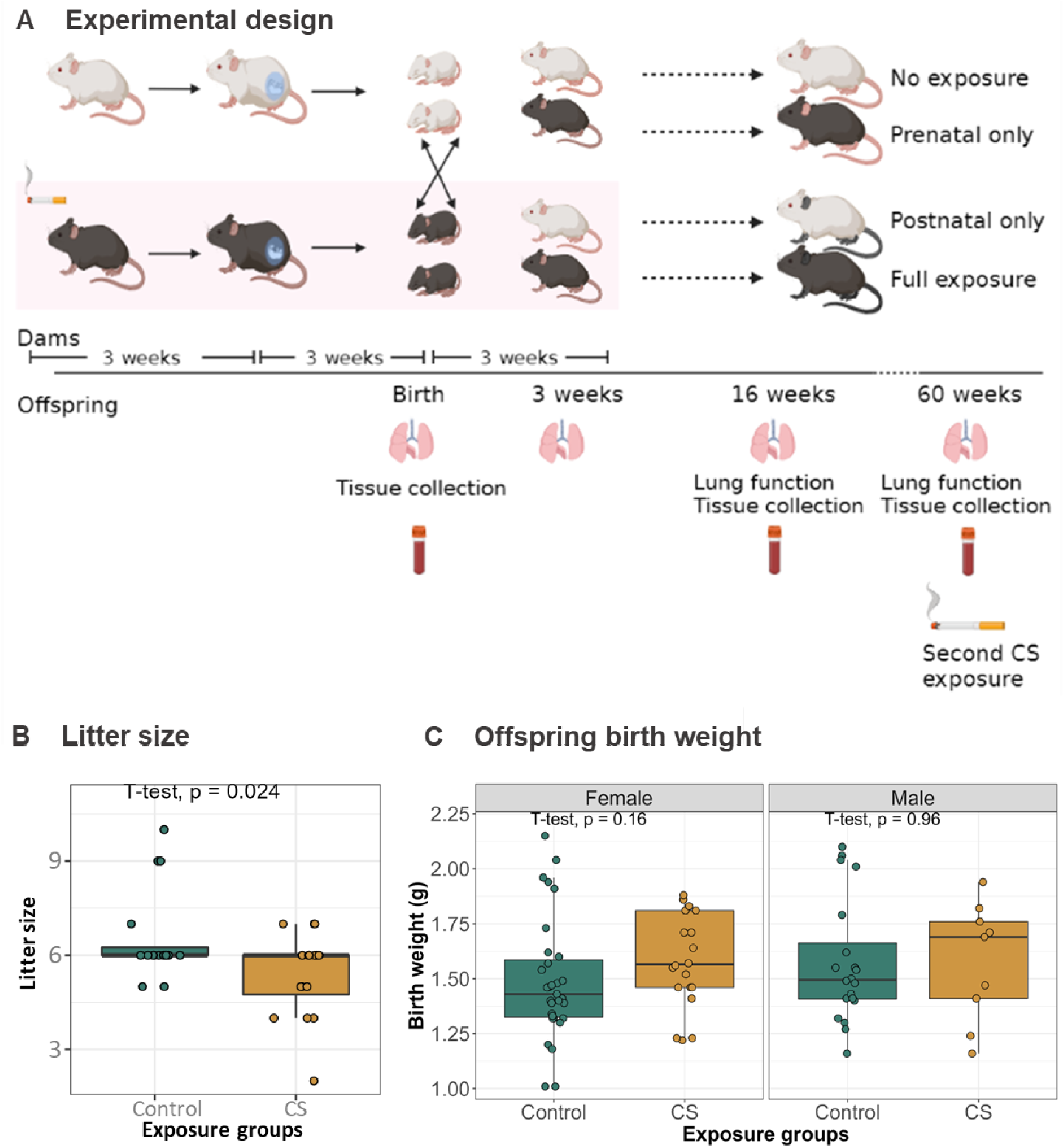
Development of a mouse model to study the effects of early life CS exposure. (A) Experimental design: Adult female mice were exposed to CS for 9 weeks, starting 3 weeks prior to mating and ending 3 weeks after birth. Half of control offspring were cross-fostered at birth with half of the CS-exposed offspring to generate 4 groups of offspring: control, prenatal CS-exposed, postnatal 13 CS-exposed and combined prenatal and postnatal CS-exposed groups. Lungs, blood and other tissues were collected from dams and offspring at birth and at 16 weeks of age, after lung function measurement. At age 60 weeks, half of the remaining female offspring were re-exposed to CS for 3 weeks, followed by lung function and tissue collection. (B) Litter size of control and smoke-exposed dams. Student t-test was used to measure differences in litter size between control and smoke-exposed dams. (C) Weight of control and CS-exposed male and female offspring at birth. Differences in birth weight were analyzed using student t-tests within sexes.

### Offspring lung phenotype at 16 weeks of age, 13 weeks after cessation of smoke exposure

At 16 weeks of age, in the absence of a methacholine challenge, male offspring with prenatal CS exposure had increased tissue elastance (Figure 2D, *p* = 0.029) and female offspring with prenatal CS exposure had increased airway resistance (Figure 2B, *p* = 0.018), compared to control offspring. When methacholine was introduced, male offspring in the postnatal CS group showed significant decrease in responsiveness for total lung resistance, airway resistance and tissue elastance (Figures 2E, 2F and 2H), while fully exposed male offspring showed increased methacholine sensitivity for total lung resistance (Figure 2E). Similarly, prenatal only and full CS exposures induced decreased methacholine responsiveness for airway resistance in female offspring (Figure 2F). While these data show increased responsiveness to specific methacholine doses in CS-exposed offspring, dose response slope analysis showed that overall, there were no significant differences in offspring sensitivity to methacholine (Supplementary Figure 1). In association with the altered lung resistance and tissue elastance, right ventricle wall thickness was increased in male and female offspring with full CS exposure (Table S4). However, female right ventricular function and hemodynamic parameters were more sensitive to the effects of CS than males (Table S5).

When examining immune cell infiltration in the lung, female offspring exposed to postnatal only CS had significantly elevated eosinophils (Mean = 1691, SD = 1647) compared to control females (Mean = 475, SD = 800) (Figure 3B, *p* = 0.0046), while male offspring with full CS exposure had significantly elevated lymphocytes (Mean = 3479, SD = 3064) compared to control males (Mean = 601, SD = 659) (Figure 3D, *p* = 0.029).

Together, our data suggested a mild lung phenotype induced by early life smoke exposure in indirectly exposed offspring that persists to 16 weeks of age, as observed by persistent decline in lung function and increased infiltration of immune cells into the lungs.

**Figure 2:**
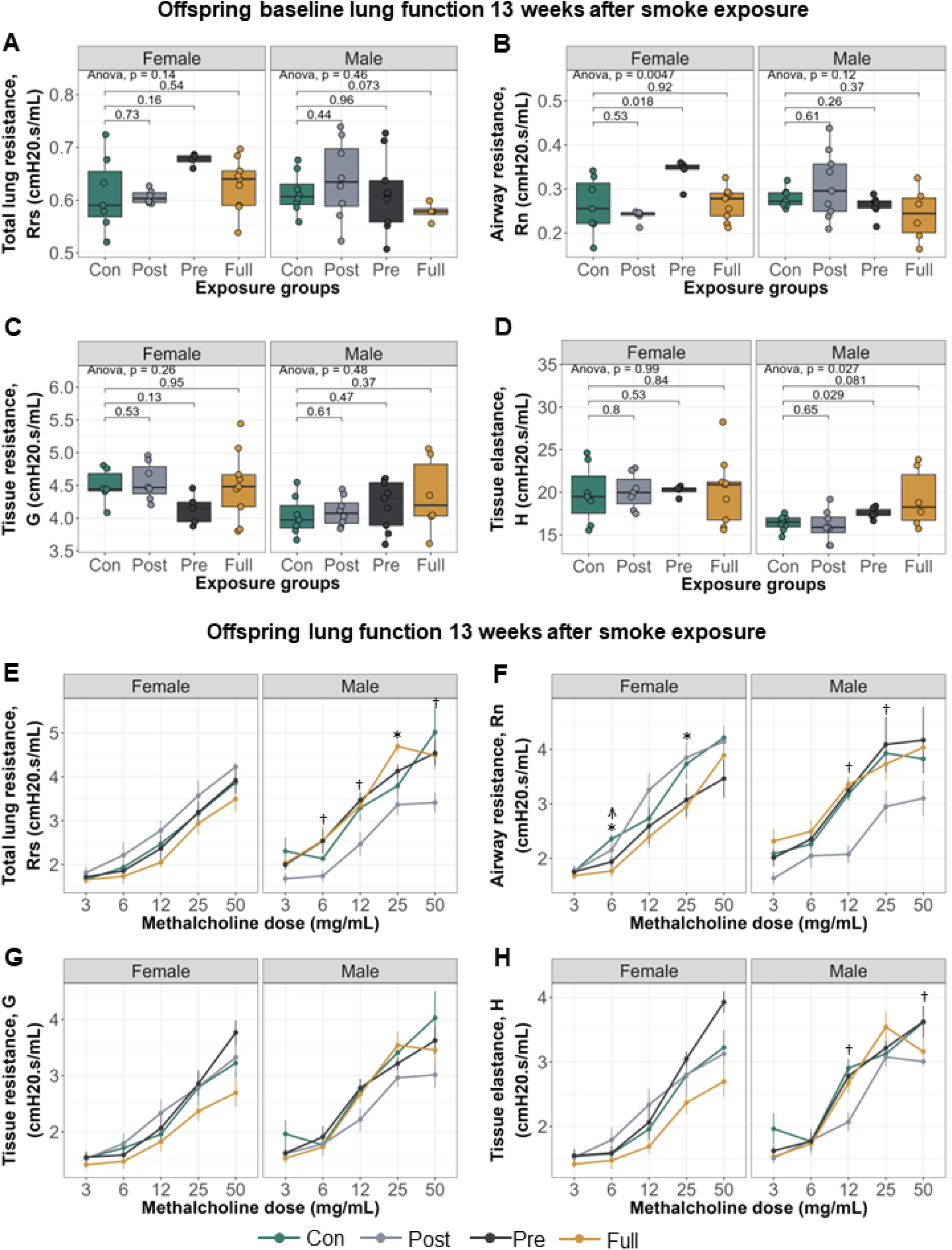
Early life exposure to CS alters sensitivity to methacholine up to 13 weeks after cessation of smoke exposure. Lung function parameters (A) Total lung resistance at baseline. (B) Airway resistance at baseline. Female offspring exposed to prenatal CS had significantly higher Rn (Mean = 0.34, SD = 0.03) compared to female controls (Mean = 0.26, SD = 0.06). (C) Tissue resistance at baseline. (D) Alveolar elastance at baseline. Male offspring exposed to prenatal CS had significantly higher H (Mean = 17.50, SD = 0.63) compared to male controls (Mean = 16.40, SD = 0.91). (E) Total lung resistance with methacholine. (F) Airway resistance with methacholine. (G) Tissue resistance with methacholine. (H) Alveolar elastance with methacholine. Lung function with methacholine (*E*–*H*): \**P* < 0.05, control vs. full CS; †*P* < 0.05 control vs postnatal CS; [r]*P* < 0.05 control vs prenatal CS groups. Lung function was assessed by taking 90^th^ percentile values in response to injection of saline into the lungs (baseline) and in response to increasing doses of methacholine. Lung function data was analyzed in a sex-disaggregated manner using a two-way ANOVA, followed by multiple comparisons between groups at each methacholine dose where significant.

**Figure 3:**
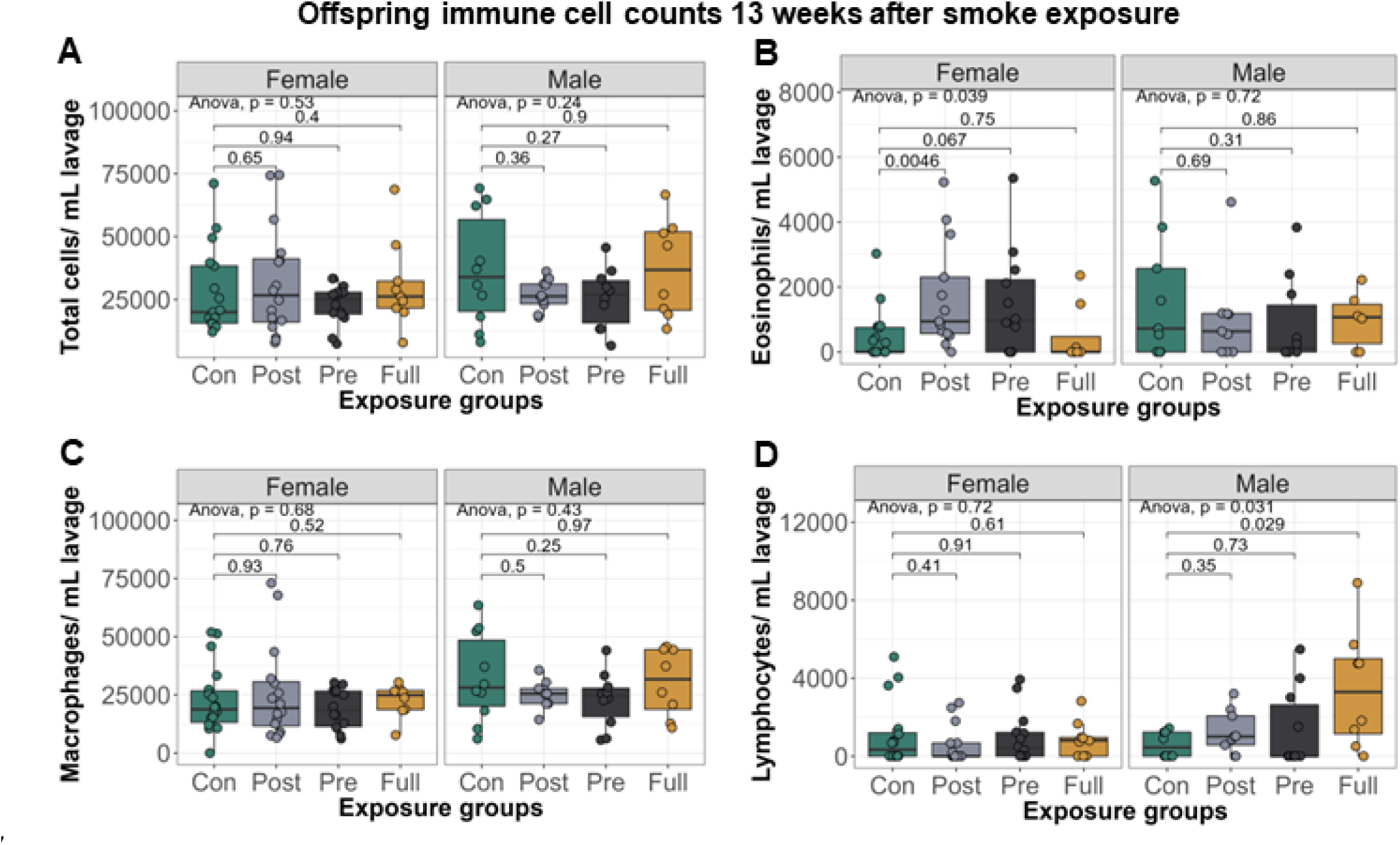
Early life exposure to CS increases immune cell infiltration into offspring lungs, up to 13 weeks after smoke exposure. (A) Total immune cells per mL lavage. (B) Eosinophils per mL lavage. (C) Macrophages per mL lavage. (D) Lymphocytes per mL lavage. Differential cell counts were normalized to lavage volume. Pairwise comparisons were conducted using t-tests.

### DNAm and gene expression at birth and 16 weeks of age in candidate genes

Results from human studies have shown that maternal smoking causes decreased and increased DNAm of *AHRR* and *CYP1A1* respectively, in newborn umbilical cord blood^19–21, 23, 28^, and that these differential methylation patterns persist in adult blood at midlife^24^. The effects of smoking on the human respiratory system is less well documented, as only one study to our knowledge has reported increased *AHRR* and *CYP1A1* DNAm in nasal epithelia of adult smokers^27^. Therefore, as DNAm is species and tissue-specific, and studies investigating effects of prenatal CS exposure on the lungs is lacking, we measured *Ahrr* and *Cyp1a1* DNAm and gene expression in the lungs and blood of offspring at birth immediately after CS exposure and at 16 weeks of age (13 weeks after cessation of smoke exposure).

At birth, *Ahrr* and *Cyp1a1* DNAm in the lung and blood were not significantly different between control and smoke-exposed offspring (Figure 4A and 4C). However, both male and female offspring born to CS-exposed dams showed lung *Cyp1a1* expression levels which were over 20 times higher than controls (Figure 4B).

We next measured *Ahrr* and *Cyp1a1* DNA methylation levels in lungs and blood of offspring at 16 weeks of age to determine longitudinal effects of early life smoke exposure on offspring (Figure 4E – 4H). There were no significant differences in *Ahrr* DNAm between control and CS-exposed groups in offspring blood 13 weeks after cessation of smoke exposure (Supplementary Figure 2). In the lung in male offspring only, *Ahrr* DNAm in prenatally exposed offspring was significantly higher than offspring exposed only postnatally (Figure 4G, *p* = 0.03221), but none of the CS-exposed offspring showed significantly altered *Ahrr* DNAm compared to control offspring. In female offspring at 16 weeks, those exposed to prenatal or postnatal CS had higher *Cyp1a1* DNAm in the lung compared to controls (Figure 4E), while offspring with full CS exposure showed significantly decreased *Cyp1a1* methylation compared to either postnatal (Figure 4E, *p* = 0.02) or prenatal groups (Figure 4E, *p* = 0.0049). These differences remained significant even after adjusting for possible effects of litter size on DNAm.

We observed the same trend when we measured offspring lung *Cyp1a1* expression: increased expression levels of *Cyp1a1* in the postnatal only and prenatal only CS groups but decreased expression in the lungs of offspring exposed to both prenatal and postnatal CS, compared to controls, though these differences were not statistically significant (Figure 4F).

To show that the differences in candidate gene DNAm observed in offspring was not because of changes in global DNAm, we measured DNAm at a control gene, *Prkaa1*, which has not been associated with smoking. As expected, *Prkaa1* methylation in lungs of offspring exposed to both prenatal and postnatal CS was not significantly different from *Prkaa1* methylation in control offspring (Supplementary Figure 3). To rule out the potential effects of cross-fostering on DNAm, we cross fostered litters without any exposures and measured *Cyp1a1* DNAm in lungs. We found no differences in *Cyp1a1* DNAm between cross-fostered and non-cross fostered offspring (Supplementary Figure 4).

This data suggested that early-life exposure to CS creates mild but long-lasting lung DNAm and gene expression changes at one of our two candidate genes.

**Figure 4:**
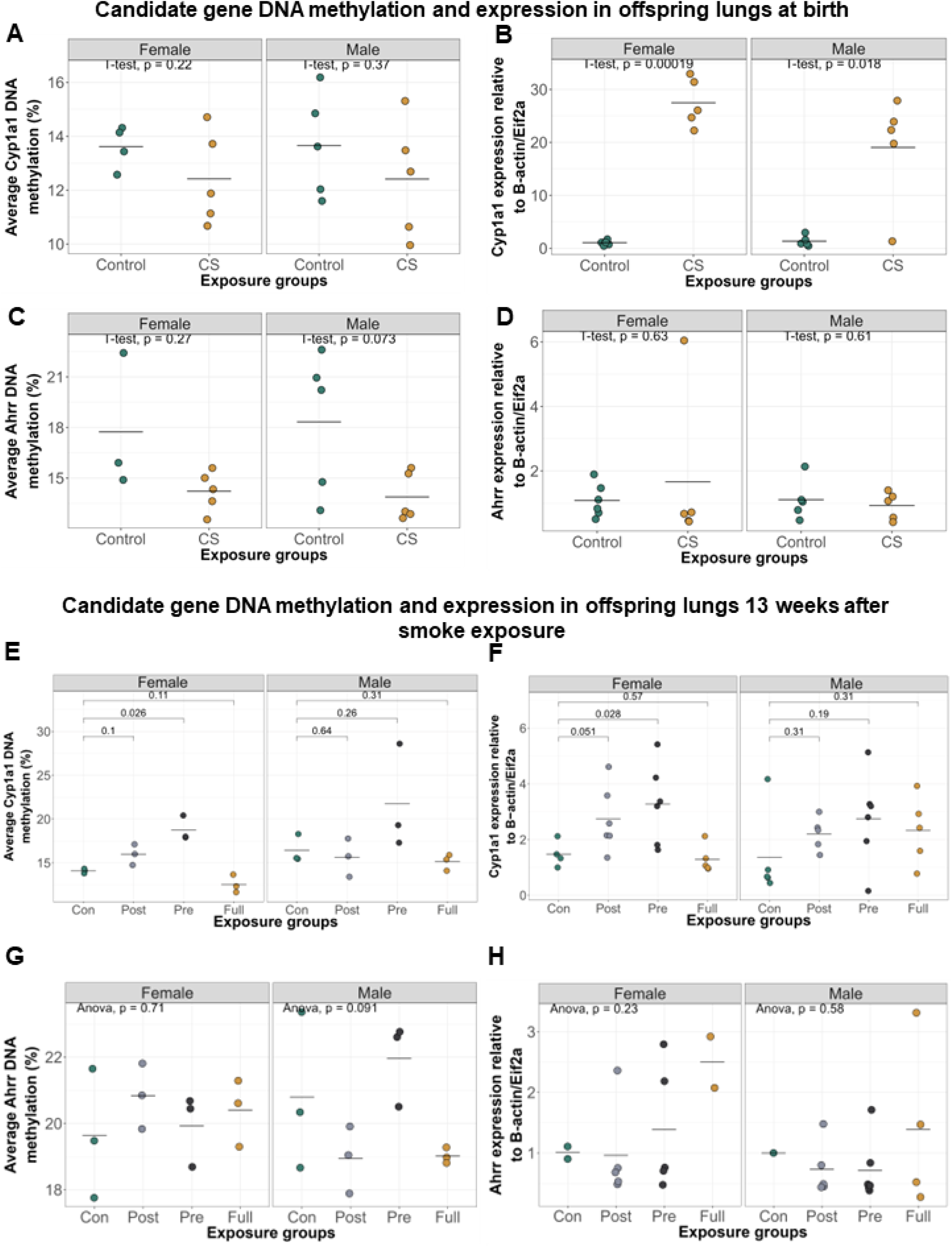
Early life exposure to CS significantly alters lung *Cyp1a1* expression at birth and *Cyp1a1* DNAm 13 weeks after smoke cessation. (A) *Cyp1a1* DNAm in offspring lungs at birth. (B) *Cyp1a1* expression in offspring lungs at birth was significantly increased in CS-exposed males (Mean = 19.00, SD = 10.30) and females (Mean = 27.50, SD = 4.54) offspring, compared to control males (Mean = 1.36, SD = 0.93) and control females (Mean = 1.47, SD = 0.43). (C) *Ahrr* DNAm in offspring lungs at birth. (D) *Ahrr* expression in offspring lungs at birth. (E) *Cyp1a1* DNAm in offspring lungs at 16 weeks of age. Prenatal and postnatal CS exposure caused an increase in female lung *Cyp1a1* DNAm, while fully exposed offspring showed a decrease. Only prenatally exposed female offspring had significantly elevated *Cyp1a1* DNAm (Mean = 18.80, SD = 1.42) compared to control females (Mean = 14.10, SD = 0.26). (F) *Cyp1a1* expression in offspring lungs at 16 weeks of age. Prenatally exposed female offspring had significantly elevated lung *Cyp1a1* expression levels (Mean = 3.28, SD = 1.44) compared to control females (Mean = 1.48, SD = 0.47). (G) *Ahrr* DNAm in offspring lungs at 16 weeks of age. (H) *Ahrr* expression in offspring lungs at 16 weeks of age. One-way ANOVA was used to compare DNAm values between groups (p<0.05 was significant), followed by two-group comparisons where significant.

### Acute re-exposure to CS at 60 weeks alters lung function and re-establishes DNAm patterns

There has been some evidence in humans that maternal smoking in pregnancy induces persistent DNAm changes until midlife, regardless of smoking habits later in life^24^. We thus sought to test what effects re-exposure to CS in adulthood would have on the lung phenotypic and DNAm patterns set by early life CS exposure. To achieve this, at 60 weeks of age, we re-exposed half of the offspring remaining to CS for 3 weeks, followed by lung function, differential immune cell analyses and DNAm measurement. At the time of re-exposure, we had a limited number of male offspring remaining, and so the following experiments were conducted solely on female offspring. We also had a low number of animals in the exclusively postnatal CS group, so they contribute to the baseline data only at 60 weeks of age.

Offspring that received full CS exposure in early life, followed by re-exposure to CS at 60 weeks, had significantly increased tissue resistance (Figure 5C) and total lung resistance (Figure 5A) levels at baseline, while re-exposed prenatal CS offspring had increased tissue resistance compared to offspring who had no early life CS exposure (controls), but were exposed to CS at 60 weeks (Figure 5). Immune cell infiltration specifically by macrophages was increased only in animals with full early life CS exposure who were also re-exposed later in life, but this increase was not statistically significant (*p* = 0.1223) (Figure 5E - 5H).

DNAm and gene expression at *Cyp1a1* showed a similar pattern, they were altered only in offspring with full CS exposure in early life who were also re-exposed to CS (Figure 6A, *p* = 0.036). This decrease in DNAm was not significant after adjusting for possible effects of litter size on DNAm using linear regression, but it is unlikely that litter size continues to influence DNAm at 60 weeks of age. Interestingly, the *Cyp1a1* DNAm patterns established upon re-exposure to CS in adulthood were similar to those observed at 16 weeks.

Taken together, this data demonstrates that early life exposure to CS induces transient changes in lung phenotype and *Cyp1a1* DNAm, which become re-established upon personal smoking in adulthood.

**Figure 5:**
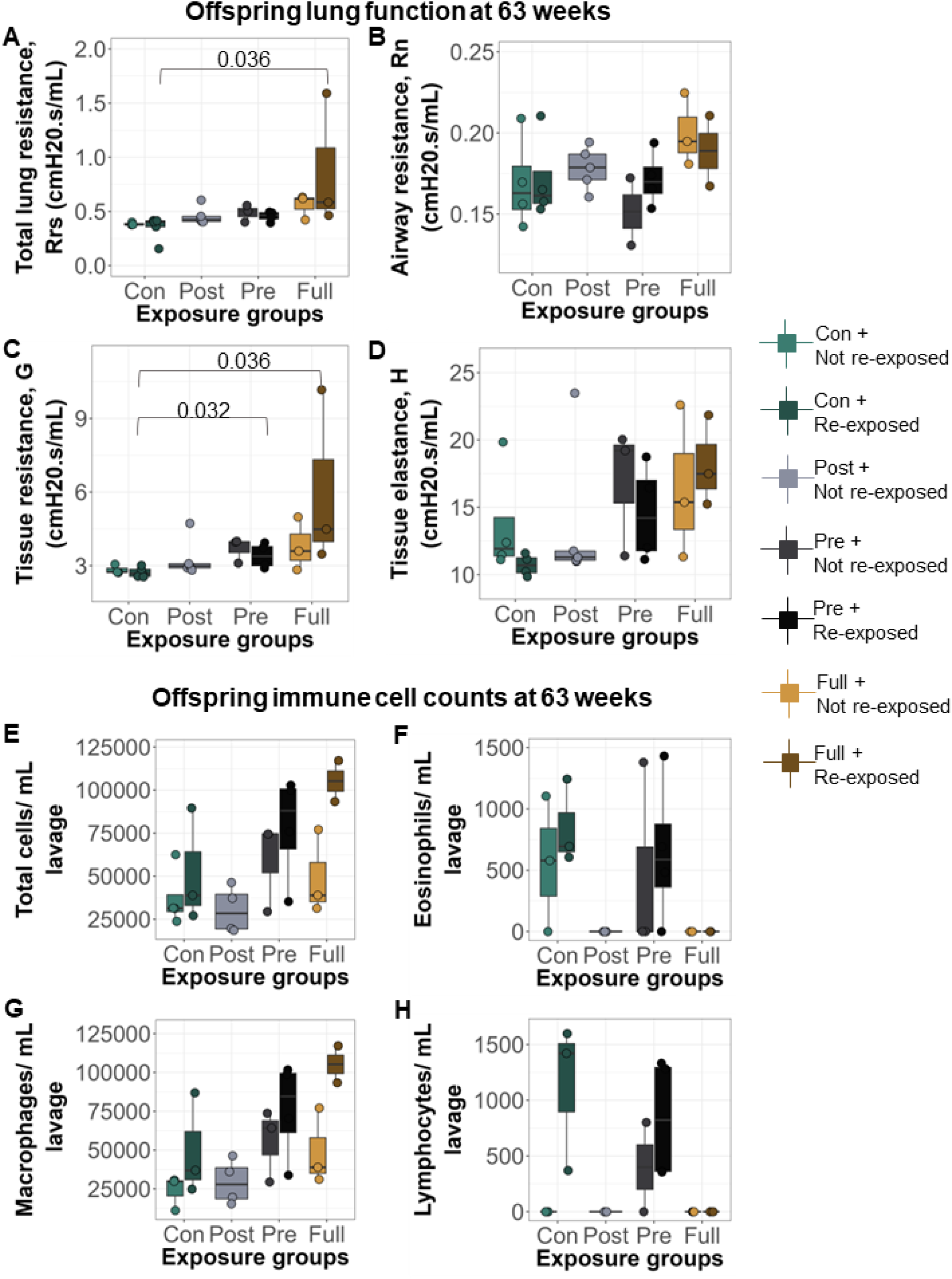
Acute re-exposure to CS at 60 weeks induced significant alterations in baseline lung function compared to early life control offspring which were exposed to CS in adulthood. Lung function parameters at baseline (A) total lung resistance. Offspring with full early life CS exposure and re-exposed to CS at 63 weeks had higher Rrs (Mean = 0.88, SD = 0.62) compared to early life control offspring exposed to CS at 63 weeks only (Mean = 0.35, SD = 0.11). (B) airway resistance. (C) tissue resistance. Offspring with full early life CS exposure and re-exposed to CS at 63 weeks had higher G (Mean = 6.04, SD = 3.60) compared to early life control offspring exposed to CS at 63 weeks only (Mean = 2.74, SD = 0.20). Offspring exposed to prenatal CS early in life and re-exposed to CS at 63 weeks had higher G (Mean = 3.40, SD = 0.51) compared to early life control offspring exposed to CS at 63 weeks only. (D) alveolar elastance. (E) Total immune cells per mL lavage. (F) Eosinophils per mL lavage. (G) Macrophages per mL lavage. (H) Lymphocytes per mL lavage. For lung function, 90^th^ percentile values were calculated in response to saline administered into the lungs. Differential cell counts were normalized to lavage volume.

**Figure 6:**
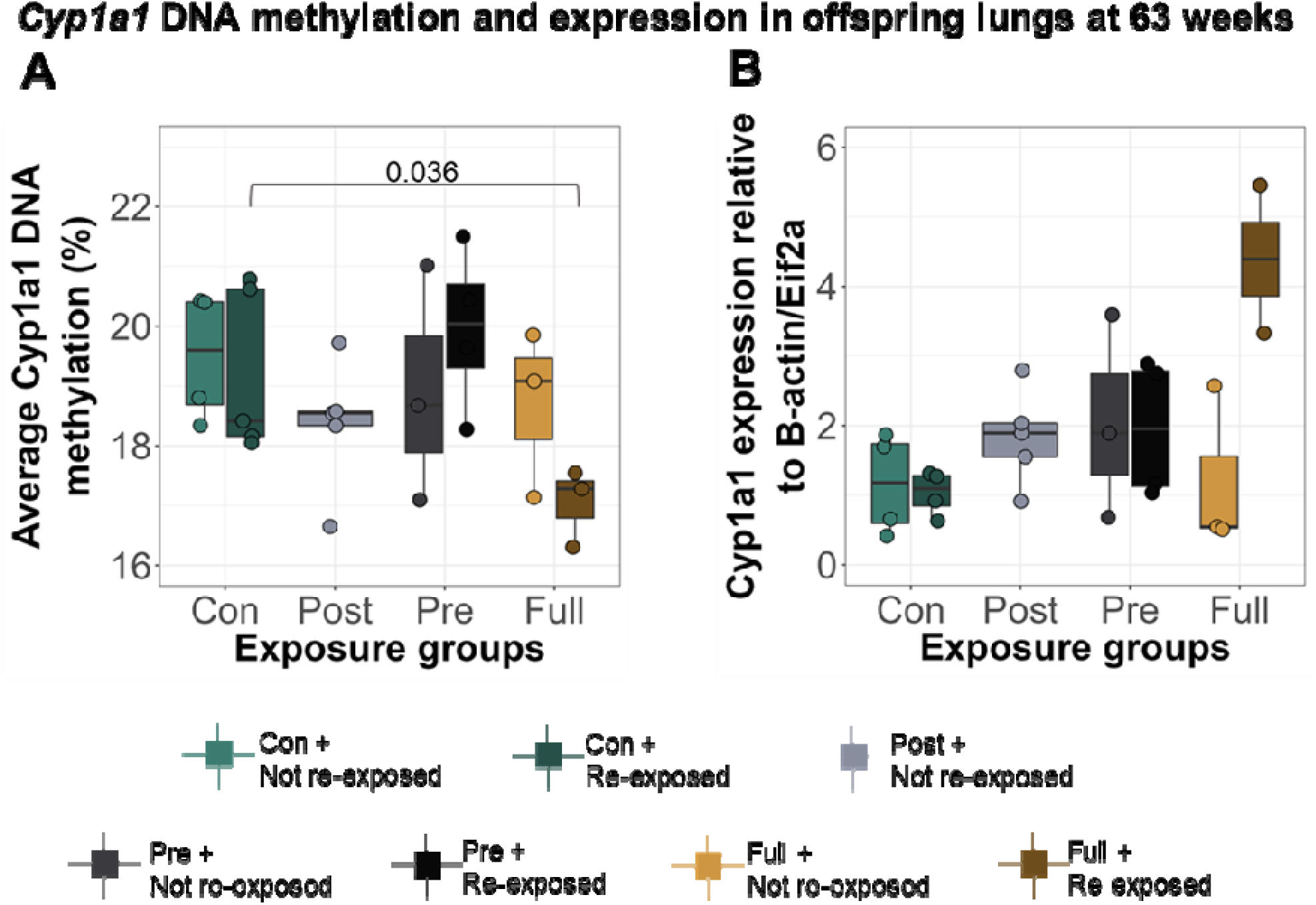
Re-exposure of offspring to CS at 60 weeks induced changes in lung DNAm which were similar to those observed at 16 weeks. (A) *Cyp1a1* DNAm in offspring lungs at 63 weeks. Offspring with full early life CS exposure and re-exposed to CS at 63 weeks had higher lung *Cyp1a1* DNAm (Mean = 17.00, SD = 0.65) compared to early life control offspring exposed to CS at 63 weeks only (Mean = 19.20, SD = 1.37). (B) *Cyp1a1* expression in offspring lungs at 63 weeks. Differences in DNAm and expression were calculated using a one-way ANOVA, followed by two-group comparisons.

## Discussion

The global burden of smoking still remains significant, despite numerous campaigns against its use, and exposure to cigarette smoke in pregnancy predisposes offspring to harmful health outcomes^48–50^. CS components have been known to pass through the placenta to the developing fetus, but it is not yet clear what specific mechanisms link CS exposure to offspring health^26, 51–53^. Past studies have shown that prenatal exposure to CS induces differential DNAm in newborn umbilical cord blood, placenta, buccal and nasal epithelia, and adult peripheral blood granulocytes^20, 23, 27, 29, 52^. However, previous studies in humans have not been able to isolate the influence of postnatal exposure to smoke, as prenatal and postnatal exposures tend to be highly correlated in humans. In addition, while the effects of maternal smoking on peripheral tissues are well documented, less is known about the effects on the lungs, despite significant epidemiological evidence linking maternal smoking to long term lung health, including asthma and COPD^54–58^. Since we know that each tissue has a unique epigenetic signature^59–67^, it is essential to measure the effects of early life smoke exposure in the lungs. Therefore, we created a mouse model to study the independent effects of prenatal and postnatal CS exposure that would also allow us to measure lung phenotype and DNAm in the lung. We used cross-fostering to isolate prenatal and postnatal exposures rather than altering exposure to CS in the dams to avoid the effects of nicotine withdrawal on the dams and pups^68–71^. Here, we showed that exposure to CS early in life alters offspring lung function, DNAm and gene expression profiles, priming the lungs such that secondary smoke exposure in adulthood re-establishes some of these outcomes. Table S3 depicts summaries of our findings.

Cigarette smoking has been shown to decrease offspring lung function^12, 56, 72–74^, but many studies identified female lungs as less sensitive to the effects of environmental toxins such as from CS^75–78^ due to earlier development of protective factors like surfactants in female lungs compared to males^79–81^. Our results largely agree with these past studies, as most of the differences in lung function we observed after methacholine exposure were in males exposed to postnatal or full CS. Only prenatally or fully exposed females showed altered airway resistance upon sensitization with methacholine. In line with past human and animal studies also^56, 78, 82^, the observed lung function defects in female offspring persisted until 16 weeks of age. For example, maternal smoking has been shown to decrease offspring lung function till 12^82^ and 21 years of age^58^. It is worth noting that while offspring showed increased *responsiveness* to different doses of methacholine at 16 weeks, overall dose response analysis showed no significant differences in methacholine *sensitivity* between groups. This implies that early life smoke exposure alone may not modulate airway reactivity but may require a secondary stimulant. To further support this finding is the fact that in female offspring, the alterations in methacholine responsiveness disappeared by 60 weeks of age but reappeared upon re-exposure to CS, suggesting that early-life CS exposure alone may not be enough to induce life-long lung function alterations but may prime the lungs so that upon re-exposure to CS, lung function is altered. This idea of priming has been well documented^77, 83, 84^, but few studies have extended exposures as long as this. One study showed that early-life secondhand smoke exposure did not alter lung function until the administration of a postnatal allergen (*A. fumigatus* extract)^84^. Another study found that *in utero* exposure to second-hand smoke exacerbated allergic response and lung function deficits in offspring when exposed to ovalbumin at 23 weeks of age^75^.

In addition to phenotype, it is important to investigate the effects of CS on lung DNAm to gain insights into the molecular mechanisms of disease development. Cytochrome P450 (Cyp) enzymes are hemoproteins which are involved in drug/xenobiotic metabolism in the liver^85^. Its major substrates are polyaromatic hydrocarbons and nitrosamines (major carcinogenic components of CS^86^), and activation by these ligands usually occurs via an aryl hydrocarbon receptor (Ahr)-dependent pathway^87, 88^. There are many subclasses of Cyp enzymes, but CYP1A1 is extensively studied as it is expressed at basal levels in extra-hepatic tissues^89^ and is highly inducible, leading to the speculation that it may be primarily responsible for xenobiotic metabolism in extra-hepatic tissues^90^. When activated by Ahr, CYP1A1 directly hydroxylates or oxidizes its bound ligand and detoxifies it^91–93^. CS is a potent inducer of the Ahr-Cyp pathway in humans and murine models^94–96^, so it follows that measuring the effects of CS on DNAm in these genes would be informative on CS-induced phenotypes.

The dynamics of *Cyp1a1* DNAm and expression over the life course in CS-exposed lungs revealed unexpected patterns. First, we noted that maternal smoking did not alter offspring *Ahrr* or *Cyp1a1* DNAm in the lung at birth at our chosen CpGs but causes a 20-fold increase in expression of just *Cyp1a1*. At 16 weeks, offspring exposed to only prenatal CS or only postnatal CS showed a slight increase in *Cyp1a1* DNAm (and expression), while those exposed to both prenatal and postnatal CS showed decreased *Cyp1a1* DNAm and expression in their left lungs compared to control offspring. We tested the effects of cross-fostering independently and saw no difference in *Cyp1a1* DNAm, which implies these differences are not an effect of cross-fostering itself. These opposing results could possibly be explained by the *developmental mismatch hypothesis*, which surmises that traits that develop in an organism in one environment may be disadvantageous in a different environment^97–99^. In other words, the mismatch between prenatal and postnatal environments in the cross-fostered offspring would confer different characteristics from offspring with matched prenatal and postnatal environments. The fact that the *Cyp1a1* DNAm in the prenatal CS only group and postnatal CS only group changes in the same direction emphasizes the need for future animal models to isolate and study them separately, as it is evident that they produce different effects from combined pre- and post-natal exposure, and can be difficult to differentiate in more long-term studies. The increase in *Cyp1a1* DNAm observed in the prenatal CS only group is in fact in line with many human studies which have measured prenatal CS-induced DNAm changes in newborn umbilical cord blood^20–23, 25^.

Finally, we observed that while early life CS-induced *Cyp1a1* DNAm changes do not persist till 60 weeks, acute exposure to CS in adulthood re-established DNAm patterns observed at 16 weeks and was associated with an immune cell infiltration phenotype that was dependent on both early life and later life CS exposure. The disappearance and re-establishment of the *Cyp1a1* DNAm patterns at 60 weeks proves that DNAm alone cannot be the epigenetic mark responsible for long-term memory of early life CS at *Cyp1a1*. The next most likely candidate for a mechanism linking early life exposure with later life response is histone modifications. One study has in fact shown that early life smoke exposure in rats causes chromatin remodeling, marked by increased histone acetylation and altered transcription factor binding in the lungs^100^. The DNAm and lung phenotype changes observed at this timepoint are further evidence of priming. Some studies have suggested that disappearance of detrimental phenotypes over time is an adaptation mechanism in response to adversity^101, 102^, which allows affected individuals to function as close to optimal as possible. However, the underlying detrimental signatures are embedded until a secondary stimulant is introduced^101, 102^.

Taken together, our results represent a novel model for the DOHaD hypothesis, which surmises that the early life environment can shape offspring phenotype to produce long term consequences in an individual. While some hypotheses such as the thrifty phenotype^103^ and adaptive response^104^ have been used to explain the longevity of health conditions following early-life adversity, the underlying mechanisms are largely unknown^105^. Epigenetic mechanisms are suspected to play major roles in this phenomenon and there are many studies which have investigated the effects of early life environment on offspring DNAm. For example, early life smoke exposure and air pollution exposure in general has been shown to cause conditions like asthma and cancer in adulthood, and many studies have suggested DNAm as the underlying mechanism^20, 31, 106, 107^. However, we have shown that in our model DNAm was not responsible for long-term CS-associated phenotype, and future work will investigate other epigenetic marks.

One limitation of this study is that we measured DNAm in whole mouse lungs and so, cannot say what specific lung cell type is responsible for the observed changes in DNAm. It is unlikely that immune cell infiltration is the cause of the DNAm changes, as lungs were lavaged to remove most of the immune cells, and the relative number of immune cells residing in lung tissue is very small. A second limitation is that our model involves administration of heavy doses of CS and so we cannot extrapolate these results to analyze the effects of light or moderate smoking.

In conclusion, this study has created an effective model for DOHaD to investigate the effects of early life CS exposure, providing an essential foundation for future animal studies. Using this model, we have shown that the effects of early life smoke exposure are embedded into the offspring, producing phenotypes which are re-established by smoking in adulthood. Future studies are needed to investigate these effects across the mouse epigenome and measure the effects of early life CS on chromatin accessibility and specific histone marks or variants, to identify the specific mechanism responsible for the long-term memory due to early life CS exposure.

## Acknowledgements

The authors thank Dr. Neeloffer Mookherjee for discussions of experimental design and helpful suggestions.

## Supplementary Information

### Methods

#### Transthoracic echocardiography

As described previously^108^, serial echocardiography was performed using a Vevo 2100 ultrasound (Fujifilm-Visual Sonics, ON Canada) equipped with a 30 MHz transducer to assess cardiac morphology and function in 12-14 week-old mice. Imaging by a blinded and experienced sonographer was performed under mild anesthesia (induced with 3% isoflurane and 1.0L/min oxygen and maintained at 1 – 1.5% isoflurane and 1.0 L/min oxygen) during echocardiography. Pulse Wave Doppler Mode was used to measure the velocity time integral of blood flow passing through the right ventricular outflow tract (RVOT). Vevo software was used to determine pulmonary acceleration times (PAT), pulmonary ejection times (PET), and ventricle thickness from the RVOT for each subject.

Measurements of PAT and PET were taken from three different doppler waves for each subject and ventricle thickness was measured from three image stills. The average of these three measurements was then recorded.

### Results

#### Impact of CS on offspring cardiac morphometry and function

To determine whether CS exposure affected cardiac morphometry or function we used echocardiography to image the heart and examine left ventricular and right ventricular structural and functional parameters. In males, there were no effects of smoke exposure on left ventricle (LV) mass or other LV structural parameters (Table S4). In females, prenatal CS exposure reduced the intraventricular septum, the LV internal diameter and LV volume (Table S5). On the other hand, postnatal CS exposure increased LV posterior wall thickness in females. Overall, these structural changes did not impact LV mass in females. Regarding LV functional parameters, in the males, prenatal CS exposure significantly increased LV ejection fraction and cardiac output, though postnatal CS and the combination of pre- and postnatal CS did not affect these parameters (Table S4). However, in females, CS exposure did not affect LV function in any of the groups (Table S5). Next, right ventricle (RV) morphometry and function was examined. In both males and females, RV wall thickness was increased in prenatal CS exposed offspring and by postnatal CS as well as the combination of prenatal and postnatal CS (Tables S4, S5). CS did not alter RV function and hemodynamics in males (Table S4). Female RV function and hemodynamic parameters were more sensitive to the effects of CS. Prenatal CS exposure increased the pulmonary vessel peak velocity and peak pressure (Table S5). In addition, prenatal CS exposure followed by postnatal CS also increased pulmonary valve peak velocity and peak pressure in females. Pulmonary acceleration time was significantly reduced by postnatal CS and the combination of prenatal and postnatal CS in females.

### Tables and figures

**Table S1:**
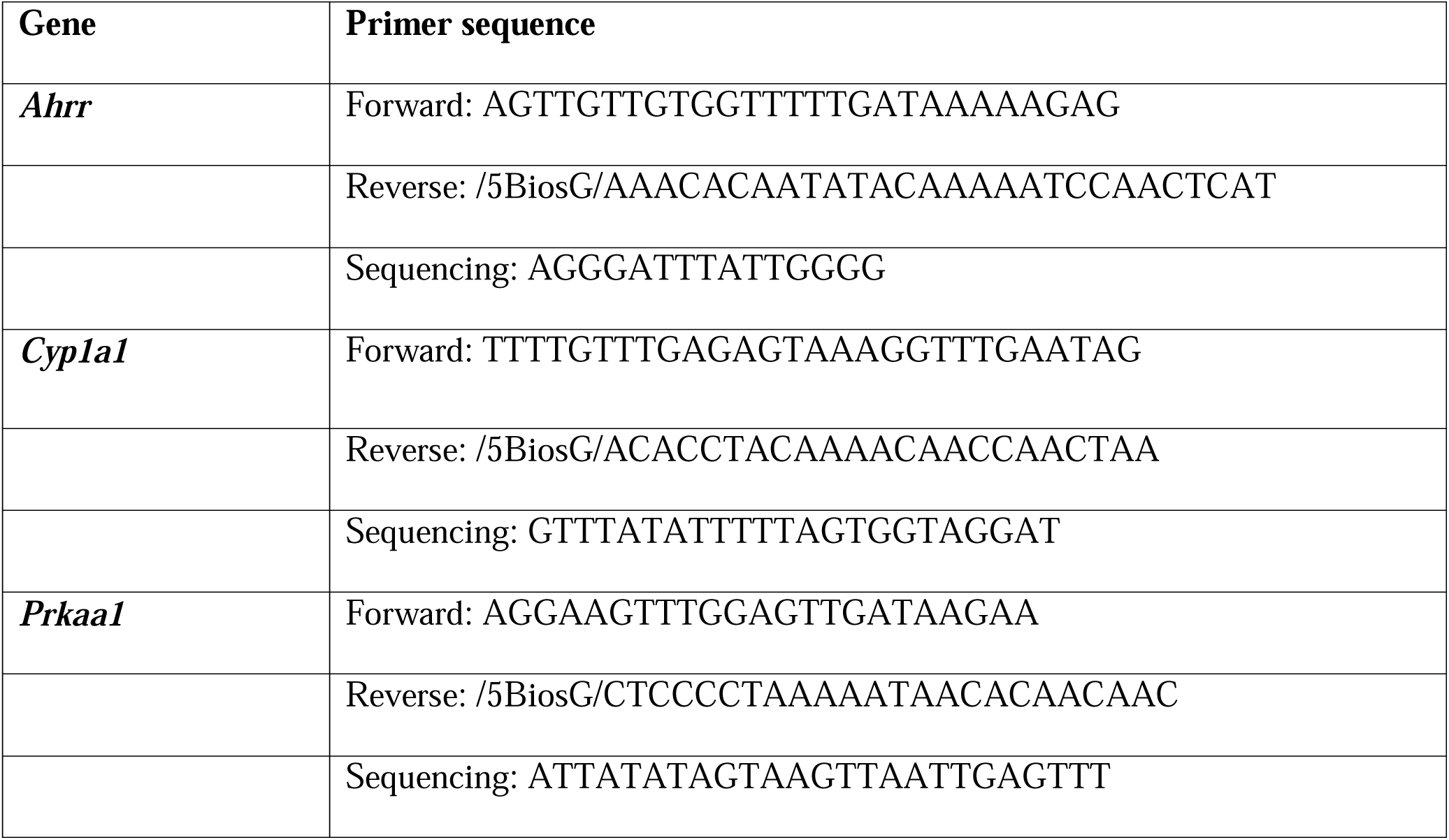
Mouse pyrosequencing primers

**Table S2:**
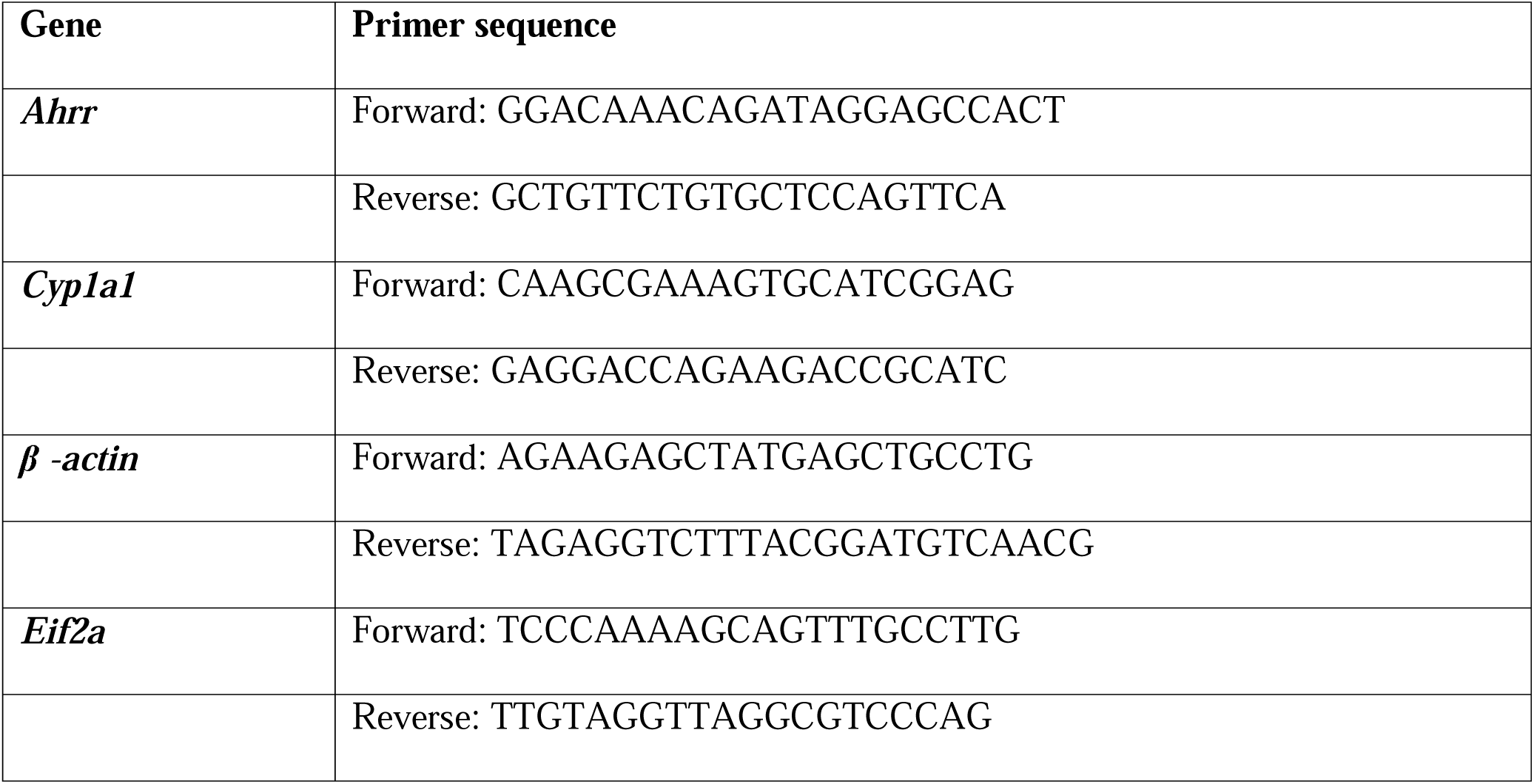
Mouse qPCR primers

**Table S3:**
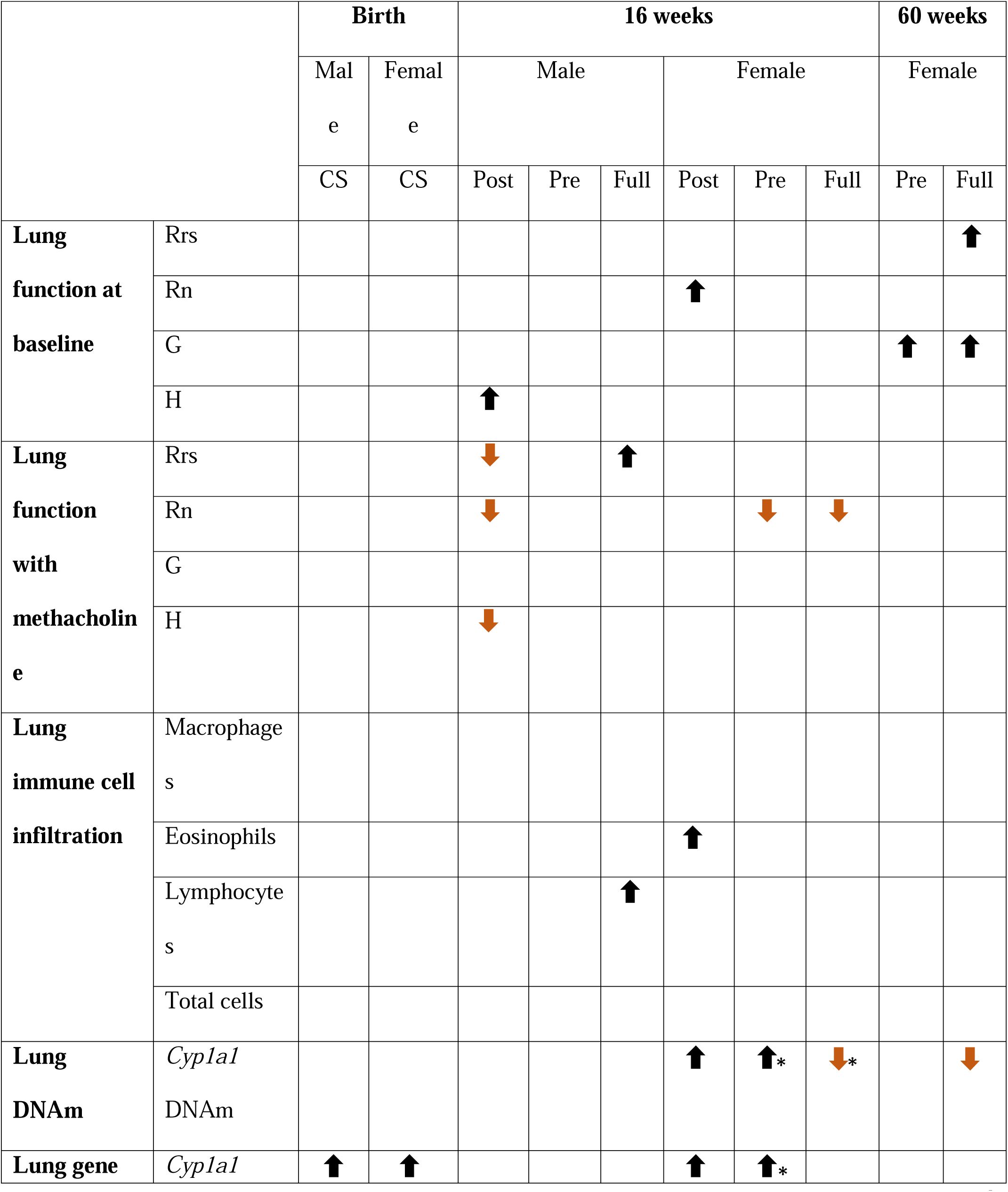
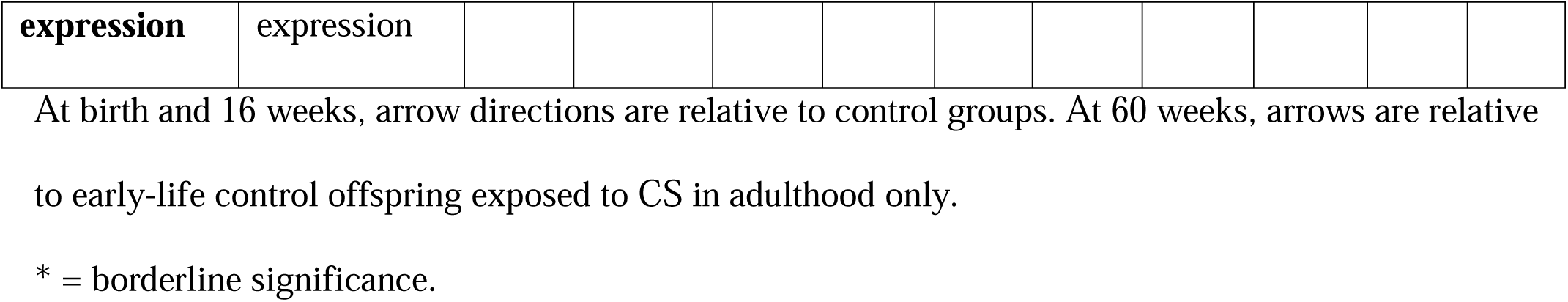
Summary of results from offspring

**Table S4:** Male offspring echocardiography parameters

|  | Control | Prenatal<br>Smoke | Postnatal<br>Smoke | Prenatal and<br>Postnatal Smoke | Prenatal<br>Smoke | Postnatal<br>Smoke | Interaction |
| --- | --- | --- | --- | --- | --- | --- | --- |
| Body<br>Weight (g) | 26.0 ±<br>0.13 | 25.9 ±<br>0.13 | 25.7 ±<br>0.06 | 25.7 ± 0.12 | 0.8872 | 0.4964 | 0.8655 |
| LV mass<br>(mg) | 89.1 ± 0.9 | 89.2 ±<br>1.4 | 85.6 ± 0.8 | 82.6 ± 1.3 | 0.6957 | 0.1840 | 0.6716 |
| LVPWd<br>(mm) | 0.75 ±<br>0.01 | 0.76 ±<br>0.01 | 0.73 ±<br>0.01 | 0.67 ± 0.01 | 0.4383 | 0.1099 | 0.3685 |
| LVAWd<br>(mm) | 0.74 ±<br>0.01 | 0.75 ±<br>0.01 | 0.79 ±<br>0.01 | 0.73 ± 0.01 | 0.4717 | 0.5564 | 0.2913 |
| IVSd<br>(mm) | 0.66 ±<br>0.01 | 0.69 ±<br>0.01 | 0.71 ±<br>0.01 | 0.68 ± 0.01 | 0.9911 | 0.5504 | 0.2704 |
| LVIDd<br>(mm) | 4.10 ±<br>0.02 | 4.06 ±<br>0.02 | 4.02 ±<br>0.01 | 4.00 ± 0.01 | 0.5881 | 0.2374 | 0.9119 |
| LV Vol<br>(ul) | 73.5 ± 0.5 | 69.8 ±<br>0.8 | 68.8 ± 0.6 | 66.6 ± 0.5 | 0.2022 | 0.0914 | 0.7522 |
| EF (%) | 51.8 ± 0.6 | 56.2 ±<br>0.7 | 53.5 ± 0.5 | 54.9 ± 0.4 | 0.0352 | 0.3285 | 0.8389 |
| FS (%) | 26.3 ± 0.4 | 29.2 ±<br>0.5 | 28.2 ± 0.3 | 29.2 ± 0.3 | 0.1095 | 0.4307 | 0.4225 |
| CO<br>(ml/min) | 14.9 ± 0.3 | 17.0 ±<br>0.2 | 15.8 ± 0.1 | 16.3 ± 0.2 | 0.0347 | 0.8675 | 0.2108 |
| SV (ul) | 38.5 ± 0.6 | 40.5 ± | 37.9 ± 0.4 | 39.5 ± 0.4 | 0.1241 | 0.6471 | 0.4787 |
|  |  | 0.5 |  |  |  |  |  |
| IVCT (ms) | 8.1 ± 0.3 | 9.9 ± 0.2 | 12.1 ± 0.3 | 11.3 ± 0.3 | 0.6817 | 0.0198 | 0.2509 |
| IVRT (ms) | 15.2 ± 0.3 | 13.7 ± 0.3 | 15.3 ± 0.2 | 14.2 ± 0.2 | 0.1118 | 0.7505 | 0.8026 |
| E/A | 2.47 ± 0.07 | 2.05 ± 0.05 | 1.88 ± 0.03 | 2.16 ± 0.05 | 0.6882 | 0.1546 | 0.0419 |
| E'/A' | 1.15 ± 0.03 | 1.12 ± 0.03 | 1.03 ± 0.02 | 1.00 ± 0.02 | 0.5589 | 0.1929 | 0.8663 |
| E/E' | 47.2 ± 0.6 | 42.8 ± 0.9 | 41.1 ± 0.5 | 40.3 ± 0.8 | 0.2331 | 0.0520 | 0.3970 |
| MPI | 0.44 ± 0.01 | 0.43 ± 0.01 | 0.51 ± 0.01 | 0.46 ± 0.01 | 0.3947 | 0.0944 | 0.4189 |
| PAT (ms) | 15.8 ± 0.2 | 15.7 ± 0.2 | 16.8 ± 0.2 | 15.7 ± 0.1 | 0.3675 | 0.4982 | 0.4769 |
| PET (ms) | 62.2 ± 0.5 | 60.3 ± 0.4 | 61.2 ± 0.3 | 59.6 ± 0.5 | 0.2517 | 0.5760 | 0.9120 |
| PAT/PET | 0.26 ± 0.01 | 0.26 ± 0.01 | 0.27 ± 0.01 | 0.26 ± 0.01 | 0.8646 | 0.2981 | 0.5046 |
| PV Peak Velocity (mmHg) | 608.0 ± 6.1 | 669.9 ± 7.7 | 637.3 ± 5.1 | 639.0 ± 3.0 | 0.1371 | 0.9716 | 0.1595 |
| PV Peak Pressure | 1.49 ± 0.03 | 1.70 ± 0.04 | 1.65 ± 0.03 | 1.64 ± 0.02 | 0.3104 | 0.6548 | 0.2682 |
| (mmHg) |  |  |  |  |  |  |  |
| RVWT | 0.50 ± | 0.52 ± | 0.53 ± | 0.57 ± 0.01 | 0.0005 | < 0.0001 | 0.2881 |
| (mm) | 0.01 | 0.01 | 0.00 |  |  |  |  |
| RVOT | 1.24 ± | 1.19 ± | 1.15 ± | 1.20 ± 0.01 | 0.9959 | 0.2501 | 0.1421 |
| (mm) | 0.01 | 0.01 | 0.01 |  |  |  |  |
| MPAP | 80.2 ± | 80.3 ± | 79.6 ± | 80.3 ± 0.08 | 0.3678 | 0.4982 | 0.4769 |
| (mmHg) | 0.13 | 0.12 | 0.11 |  |  |  |  |
| HR (bpm) | 390 ± 4.1 | 405 ± | 413 ± 2.6 | 416 ± 3.6 | 0.7402 | 0.9087 | 0.2510 |
|  |  | 3.8 |  |  |  |  |  |

Abbreviations: HR-heart rate; LV- left ventricle; LVID- LV internal diameter; LVAW- LV anterior wall; LVPW- LV posterior wall; IVS- interventricular septum; EF- ejection fraction; FS- fractional shortening; CO- cardiac output; SV- stroke volume; IVRT- isovolumic relaxation time; E/A; ratio of mitral valve peak velocities measured by pulsed wave doppler during the early (E) and atrial (A) peaks; E'/A' - ratio of mitral valve peak velocities measured by tissue doppler during the early (E') and atrial (A') peaks; E/E' - ratio of early mitral inflow velocity (E) and mitral annular early diastolic velocity (E') measured by tissue doppler; MPI- myocardial performance index; LV Vol- end diastolic volume; bpm- beats per minute; PAT- pulmonary acceleration time; PET- pulmonary ejection time; RVWT- right ventricle wall thickness; RVOT- right ventricular outflow tract; MPAP- Mean pulmonary arterial pressure

**Table S5:** Female offspring echocardiography parameters

|  | Control | Prenatal<br>Smoke | Postnatal<br>Smoke | Prenatal and<br>Postnatal Smoke | Prenatal<br>Smoke | Postnatal<br>Smoke | Interaction |
| --- | --- | --- | --- | --- | --- | --- | --- |
| Body<br>Weight (g) | 21.4 ±<br>0.04 | 21.3 ±<br>0.07 | 21.1 ±<br>0.06 | 20.5 ± 0.07 | 0.2274 | 0.0658 | 0.4613 |
| LV mass<br>(mg) | 68.1 ± 0.3 | 67.5 ±<br>0.5 | 69.0 ± 0.5 | 68.0 ± 0.6 | 0.7459 | 0.6094 | 0.9065 |
| LVPWd<br>(mm) | 0.61 ±<br>0.002 | 0.63 ±<br>0.003 | 0.66 ±<br>0.005 | 0.69 ± 0.005 | 0.1866 | 0.0010 | 0.9604 |
| LVAWd<br>(mm) | 0.65 ±<br>0.003 | 0.61 ±<br>0.003 | 0.65 ±<br>0.003 | 0.67 ± 0.005 | 0.5532 | 0.0733 | 0.1731 |
| IVSd<br>(mm) | 0.65 ±<br>0.003 | 0.62 ±<br>0.003 | 0.67 ±<br>0.003 | 0.65 ± 0.004 | 0.05 | 0.1592 | 0.6054 |
| LVIDd<br>(mm) | 3.7 ± 0.01 | 3.6 ±<br>0.01 | 3.7 ± 0.01 | 3.6 ± 0.01 | 0.0072 | 0.5422 | 0.3386 |
| LV Vol<br>(ul) | 57.8 ±<br>0.15 | 55.9 ±<br>0.28 | 58.9 ±<br>0.34 | 53.9 ± 0.38 | 0.0112 | 0.7292 | 0.2413 |
| EF (%) | 57.1 ± 0.2 | 57.0 ±<br>0.4 | 55.3 ± 0.3 | 59.2 ± 0.3 | 0.1831 | 0.8515 | 0.1593 |
| FS (%) | 29.0 ± 0.1 | 28.7 ±<br>0.2 | 28.4 ± 0.2 | 30.9 ± 0.2 | 0.2157 | 0.3742 | 0.1142 |
| CO<br>(ml/min) | 13.4 ±<br>0.07 | 12.5 ±<br>0.09 | 13.2 ±<br>0.09 | 13.2 ± 0.11 | 0.3023 | 0.5938 | 0.2920 |
| SV (ul) | 34.1 ± | 32.9 ± | 33.3 ± | 33.2 ± 0.24 | 0.5342 | 0.8299 | 0.6036 |
|  | 0.13 | 0.25 | 0.24 |  |  |  |  |
| IVCT (ms) | 10.6 ±<br>0.14 | 10.3 ±<br>0.16 | 11.4 ±<br>0.15 | 9.4 ± 0.18 | 0.1162 | 0.9182 | 0.2574 |
| IVRT (ms) | 13.9 ±<br>0.09 | 14.8 ±<br>0.14 | 13.4 ±<br>0.10 | 14.0 ± 0.12 | 0.1287 | 0.1935 | 0.7663 |
| E/A | 2.44 ±<br>0.02 | 2.37 ±<br>0.03 | 2.38 ±<br>0.03 | 2.07 ± 0.03 | 0.2039 | 0.2141 | 0.2858 |
| E'/A' | 1.04 ±<br>0.01 | 1.08 ±<br>0.01 | 1.05 ±<br>0.01 | 1.11 ± 0.01 | 0.3278 | 0.6450 | 0.8636 |
| E/E' | 41.0 ±<br>0.31 | 39.9 ±<br>0.40 | 41.2 ±<br>0.33 | 39.5 ± 0.40 | 0.4263 | 0.9619 | 0.8685 |
| MPI | 0.44 ±<br>0.003 | 0.44 ±<br>0.004 | 0.45 ±<br>0.004 | 0.44 ± 0.004 | 0.6185 | 0.9261 | 0.9677 |
| PAT (ms) | 16.9 ±<br>0.10 | 15.9 ±<br>0.08 | 15.5 ±<br>0.07 | 15.0 ± 0.08 | 0.0888 | 0.0105 | 0.4307 |
| PET (ms) | 63.0 ±<br>0.14 | 64.2 ±<br>0.17 | 62.1 ±<br>0.22 | 62.0 ± 0.17 | 0.5380 | 0.0826 | 0.4368 |
| PAT/PET | 0.27 ±<br>0.002 | 0.25 ±<br>0.001 | 0.25 ±<br>0.001 | 0.24 ± 0.001 | 0.0903 | 0.0894 | 0.3316 |
| PV Peak<br>Vel<br>(mm/s) | 580.0 ±<br>2.3 | 615.1 ±<br>3.9 | 605.6 ±<br>2.7 | 635.6 ± 3.4 | 0.0294 | 0.1213 | 0.8618 |
| PV Peak | 1.36 ± | 1.58 ± | 1.53 ± | 1.69 ± 0.02 | 0.0080 | 0.0494 | 0.7303 |
| Pressure<br>(mmHg) | 0.01 | 0.02 | 0.01 |  |  |  |  |
| RVWT<br>(mm) | 0.46 ±<br>0.000 | 0.49 ±<br>0.002 | 0.51 ±<br>0.003 | 0.54 ± 0.002 | 0.0004 | < 0.0001 | 0.2874 |
| RVOT<br>(mm) | 1.08 ±<br>0.004 | 1.13 ±<br>0.004 | 1.10 ±<br>0.004 | 1.14 ± 0.005 | 0.0609 | 0.4914 | 0.5624 |
| MPAP<br>(mmHg) | 79.5 ±<br>0.06 | 79.7 ±<br>0.07 | 80.1 ±<br>0.05 | 80.5 ± 0.06 | 0.4103 | 0.0265 | 0.7468 |
| HR (bpm) | 393 ± 1.1 | 389 ±<br>1.5 | 398 ± 1.6 | 397 ± 1.3 | 0.7349 | 0.3194 | 0.8034 |

Abbreviations: HR-heart rate; LV- left ventricle; LVID- LV internal diameter; LVAW- LV anterior wall; LVPW- LV posterior wall; IVS- interventricular septum; EF- ejection fraction; FS- fractional shortening; CO- cardiac output; SV- stroke volume; IVRT- isovolumic relaxation time; E/A; ratio of mitral valve peak velocities measured by pulsed wave doppler during the early (E) and atrial (A) peaks; E'/A'- ratio of mitral valve peak velocities measured by tissue doppler during the early (E') and atrial (A') peaks; E/E'- ratio of early mitral inflow velocity (E) and mitral annular early diastolic velocity (E') measured by tissue doppler; MPI- myocardial performance index; LV Vol- end diastolic volume; bpm- beats per minute; PAT- pulmonary acceleration time; PET- pulmonary ejection time; RVWT- right ventricle wall thickness; RVOT- right ventricular outflow tract; MPAP- Mean pulmonary arterial pressure

**Supplementary Figure 1:**
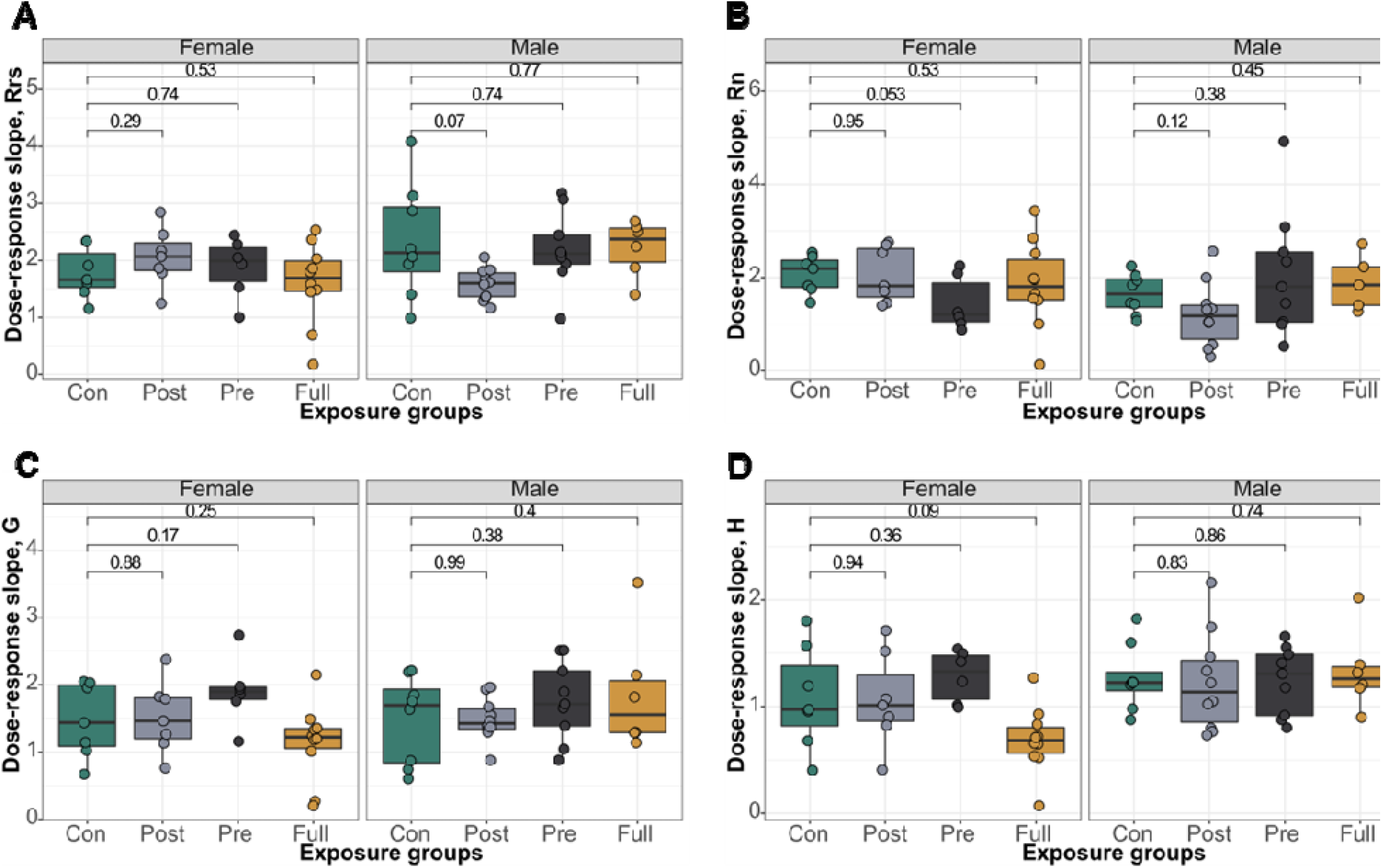
Methacholine dose response slope analyses for lung function measurements in male and female offspring at 16 weeks. A) Dose-response slope calculation for total lung resistance B) Dose-response slope for airway resistance C) Dose-response slope calculation for tissue resistance D) Dose-response slope calculation for alveolar elastance. p<0.05 (t-test) was significant.

**Supplementary Figure 2:**
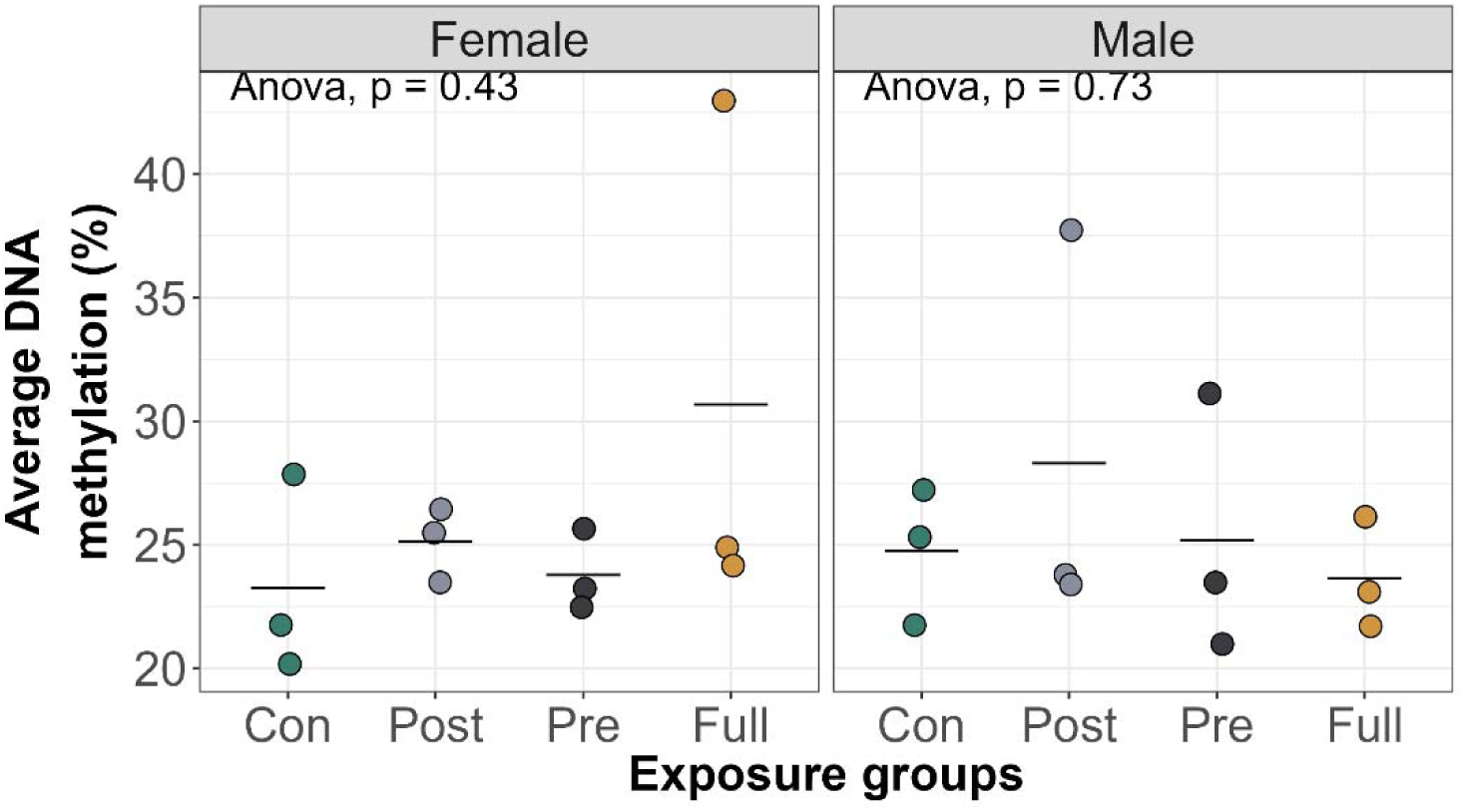
Smoking did not alter *Ahrr* DNAm in offspring blood 13 weeks after smoke cessation.

**Supplementary Figure 3:**
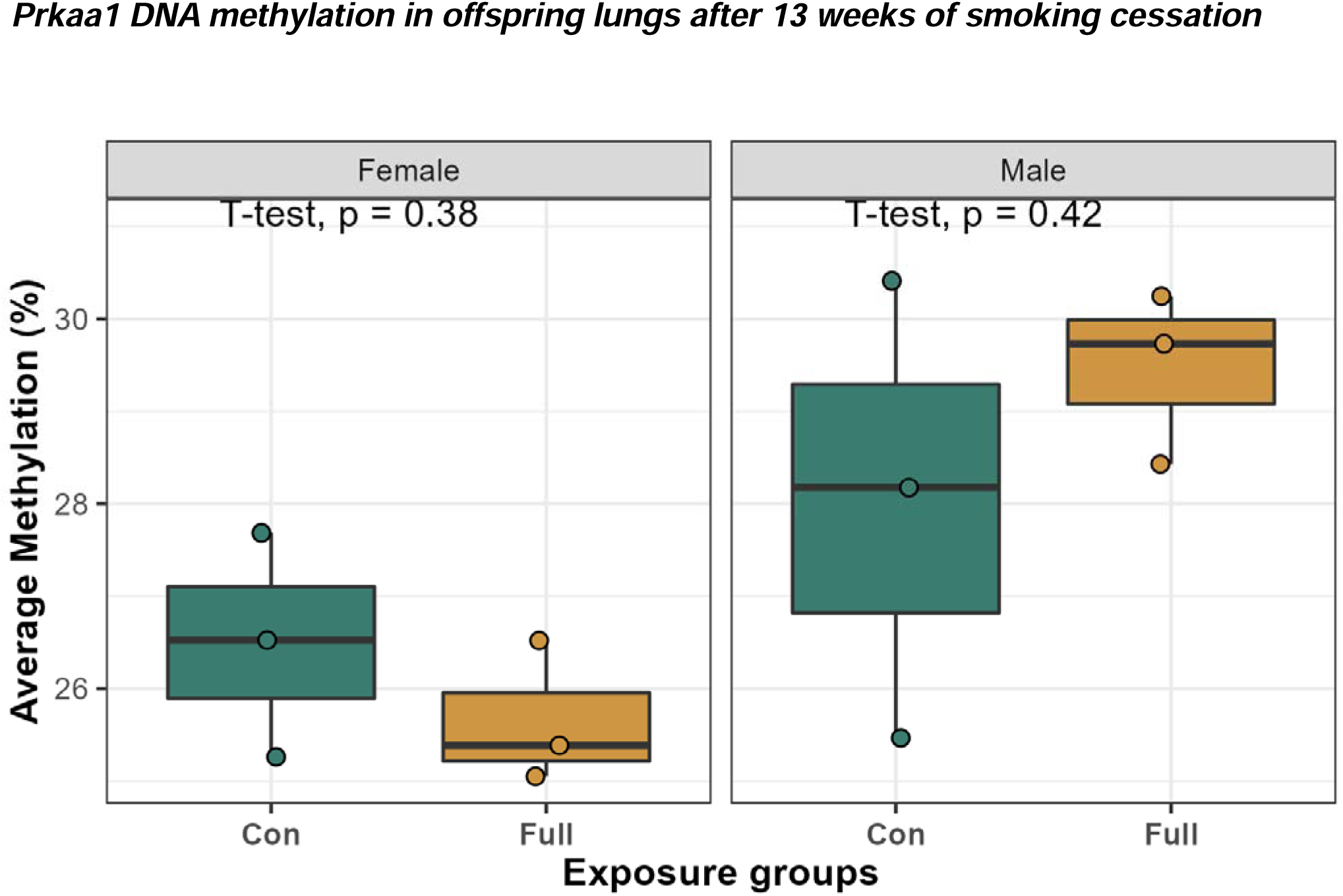
Early life exposure to CS did not significantly alter offspring lung DNAm at the control gene, *Prkaa1*.

**Supplementary Figure 4:**
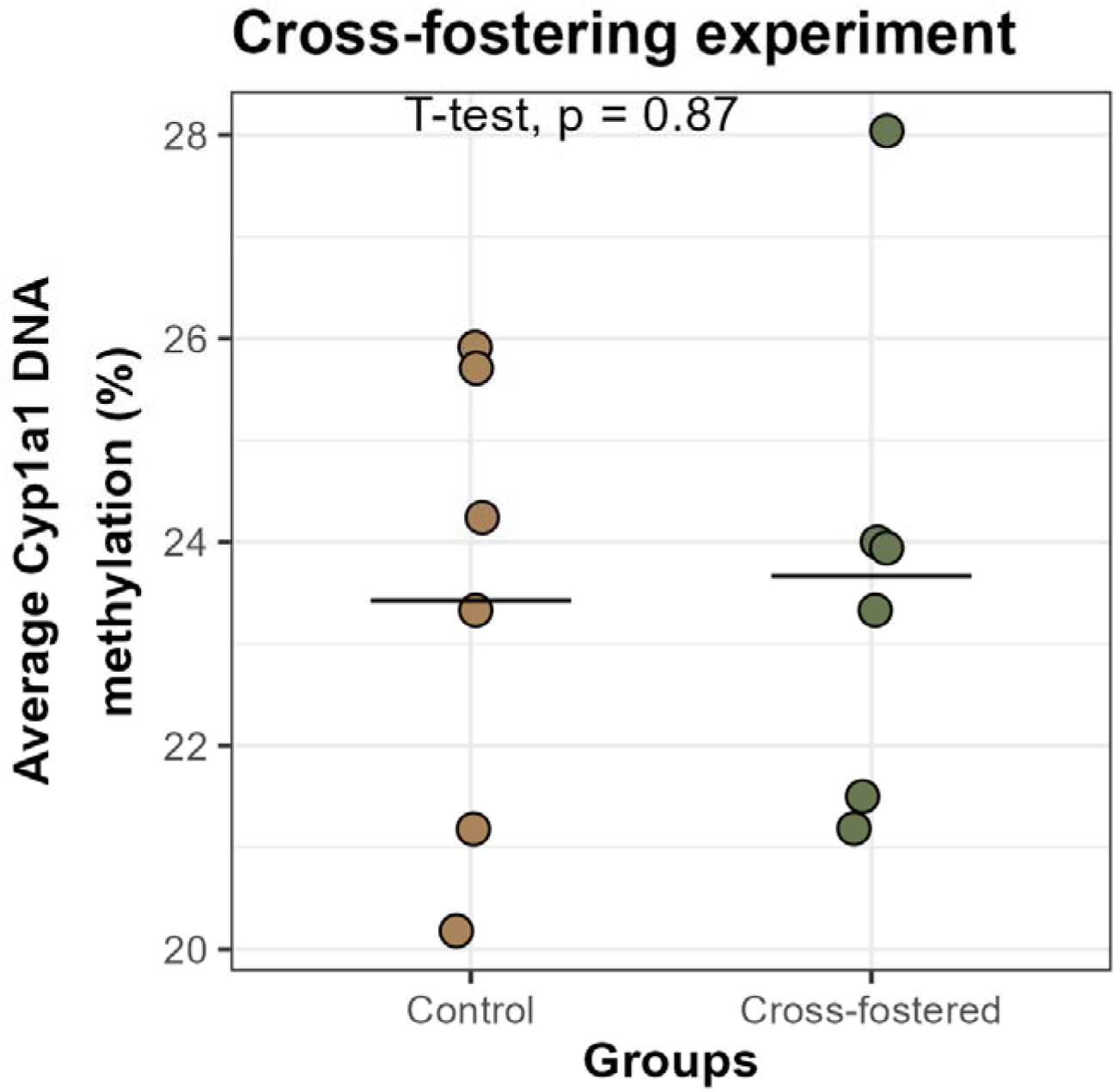
Cross-fostering did not significantly affect *Cyp1a1* DNAm in mouse lungs. To rule out potential effects of cross-fostering on offspring DNAm, we measured *Cyp1a1* DNAm in lungs of cross-fostered and non-cross fostered control Balb/C mice at 8 weeks of age. Differences in DNAm were measured using a student t-test.

## References

1. Roseboom, T. J. et al. Effects of Prenatal Exposure to the Dutch Famine on Adult Disease in Later Life: An Overview. Twin Research and Human Genetics 4, 293–298 (2001).

2. Nielsen, C. H., Larsen, A. & Nielsen, A. L. DNA methylation alterations in response to prenatal exposure of maternal cigarette smoking: A persistent epigenetic impact on health from maternal lifestyle? Arch Toxicol 90, 231–245 (2016).

3. Gluckman, P. D., Hanson, M. A., Cooper, C. & Thornburg, K. L. Effect of in utero and early-life conditions on adult health and disease. The New England journal of medicine 359, 61–73 (2008).

4. Mandy, M. & Nyirenda, M. Developmental Origins of Health and Disease: the relevance to developing nations. International Health 10, 66–70 (2018).

5. Government of Canada, S. C. Smokers, by age group. https://www150.statcan.gc.ca/t1/tbl1/en/tv.action?pid=1310009610 (2018).

6. Harris, J. E. Cigarette Smoke Components and Disease: Cigarette Smoke I s More Than a Triad of Tar, Nicotine, and Carbon Monoxide. 17.

7. Swan, G. E. & Lessov-Schlaggar, C. N. The Effects of Tobacco Smoke and Nicotine on Cognition and the Brain. Neuropsychol Rev 17, 259–273 (2007).

8. Cheraghi, M. & Salvi, S. Environmental tobacco smoke (ETS) and respiratory health in children. Eur J Pediatr 168, 897–905 (2009).

9. Ekblad, M., Korkeila, J. & Lehtonen, L. Smoking during pregnancy affects foetal brain development. Acta Paediatrica 104, 12–18 (2015).

10. Rogers, J. M. Tobacco and pregnancy. Reproductive Toxicology 28, 152–160 (2009).

11. CDCTobaccoFree. 2014 SGR: The Health Consequences of Smoking—50 Years of Progress. Centers for Disease Control and Prevention https://www.cdc.gov/tobacco/data_statistics/sgr/50th-anniversary/index.htm (2021).

12. Singh, S. P. et al. Prenatal Cigarette Smoke Decreases Lung cAMP and Increases Airway Hyperresponsiveness. Am J Respir Crit Care Med 168, 342–347 (2003).

13. Lawder, R., Whyte, B., Wood, R., Fischbacher, C. & Tappin, D. M. Impact of maternal smoking on early childhood health: a retrospective cohort linked dataset analysis of 697 003 children born in Scotland 1997–2009. BMJ Open 9, (2019).

14. Hertzman, C. & Wiens, M. Child development and long-term outcomes: a population health perspective and summary of successful interventions. Soc Sci Med 43, 1083–1095 (1996).

15. Aristizabal, M. J. et al. Biological embedding of experience: A primer on epigenetics. PNAS (2019) doi:10.1073/pnas.1820838116.

16. Demetriou, C. A. et al. Biological embedding of early-life exposures and disease risk in humans: a role for DNA methylation. Eur. J. Clin. Invest. 45, 303–332 (2015).

17. Cunliffe, V. T. Experience-sensitive epigenetic mechanisms, developmental plasticity, and the biological embedding of chronic disease risk. WIREs Systems Biology and Medicine 7, 53–71 (2015).

18. Kim, J. K., Samaranayake, M. & Pradhan, S. Epigenetic mechanisms in mammals. Cell. Mol. Life Sci. 66, 596 (2008).

19. Joubert, B. R. et al. 450K epigenome-wide scan identifies differential DNA methylation in newborns related to maternal smoking during pregnancy. Environmental health perspectives 120, 1425–1431 (2012).

20. Joubert, B. R. et al. DNA Methylation in Newborns and Maternal Smoking in Pregnancy: Genome-wide Consortium Meta-analysis. American journal of human genetics 98, 680–696 (2016).

21. Richmond, R. C. et al. Prenatal exposure to maternal smoking and offspring DNA methylation across the lifecourse: findings from the Avon Longitudinal Study of Parents and Children (ALSPAC). Hum Mol Genet 24, 2201–2217 (2015).

22. Wiklund, P. et al. DNA methylation links prenatal smoking exposure to later life health outcomes in offspring. Clinical Epigenetics 11, 97 (2019).

23. Lee, K. W. K. et al. Prenatal Exposure to Maternal Cigarette Smoking and DNA Methylation: Epigenome-Wide Association in a Discovery Sample of Adolescents and Replication in an Independent Cohort at Birth through 17 Years of Age. Environ Health Perspect 123, 193–199 (2015).

24. Tehranifar, P. et al. Maternal cigarette smoking during pregnancy and offspring DNA methylation in midlife. EpigeneticsflJ: official journal of the DNA Methylation Society 13, 129–134 (2018).

25. Richmond, R. C., Suderman, M., Langdon, R., Relton, C. L. & Davey Smith, G. DNA methylation as a marker for prenatal smoke exposure in adults. Int J Epidemiol 47, 1120–1130 (2018).

26. Suter, M. et al. IN UTERO TOBACCO EXPOSURE EPIGENETICALLY MODIFIES PLACENTAL CYP1A1 EXPRESSION. Metabolism 59, 1481–1490 (2010).

27. Reddy, K. D. et al. Current Smoking Alters Gene Expression and DNA Methylation in the Nasal Epithelium of Patients with Asthma. Am J Respir Cell Mol Biol 65, 366–377 (2021).

28. Rauschert, S. et al. Maternal Smoking During Pregnancy Induces Persistent Epigenetic Changes Into Adolescence, Independent of Postnatal Smoke Exposure and Is Associated With Cardiometabolic Risk. Frontiers in Genetics 10, (2019).

29. Breton, C. V. et al. Prenatal tobacco smoke exposure affects global and gene-specific DNA methylation. Am J Respir Crit Care Med 180, 462–467 (2009).

30. Flom, J. D. et al. Prenatal smoke exposure and genomic DNA methylation in a multiethnic birth cohort. Cancer Epidemiol Biomarkers Prev 20, 2518–2523 (2011).

31. Neophytou, A. M. et al. In utero tobacco smoke exposure, DNA methylation, and asthma in Latino children. Environmental Epidemiology 3, e048 (2019).

32. Sherrill, D. L. et al. Longitudinal effects of passive smoking on pulmonary function in New Zealand children. Am Rev Respir Dis 145, 1136–1141 (1992).

33. Knopik, V. S. et al. Contributions of parental alcoholism, prenatal substance exposure, and genetic transmission to child ADHD risk: a female twin study. Psychol. Med. 35, 625–635 (2005).

34. Agrawal, A. et al. Correlates of cigarette smoking during pregnancy and its genetic and environmental overlap with nicotine dependence. Nicotine Tob Res 10, 567–578 (2008).

35. Vardavas, C. I. et al. The independent role of prenatal and postnatal exposure to active and passive smoking on the development of early wheeze in children. The European respiratory journal 48, 115–124 (2016).

36. Joad, J. P., Ji, C. M., Kott, K. S., Bric, J. M. & Pinkerton, K. E. In Utero and Postnatal Effects of Sidestream Cigarette Smoke Exposure on Lung Function, Hyperresponsiveness, and Neuroendocrine Cells in Rats. Toxicology and Applied Pharmacology 132, 63–71 (1995).

37. Burke, H. et al. Prenatal and passive smoke exposure and incidence of asthma and wheeze: systematic review and meta-analysis. Pediatrics 129, 735–744 (2012).

38. Eskenazi, B. & Castorina, R. Association of prenatal maternal or postnatal child environmental tobacco smoke exposure and neurodevelopmental and behavioral problems in children. Environmental Health Perspectives 107, 991–1000 (1999).

39. Ma, Y. et al. Sustained suppression of IL-13 by a vaccine attenuates airway inflammation and remodeling in mice. Am J Respir Cell Mol Biol 48, 540–549 (2013).

40. Ryu, M. H. et al. Chronic exposure to perfluorinated compounds: Impact on airway hyperresponsiveness and inflammation. Am J Physiol Lung Cell Mol Physiol 307, L765–774 (2014).

41. Pascoe, C. D. et al. Allergen inhalation generates pro-inflammatory oxidised phosphatidylcholine associated with airway dysfunction. Eur Respir J 57, 2000839 (2021).

42. MUSCLE: multiple sequence alignment with improved accuracy and speed | IEEE Conference Publication | IEEE Xplore. https://ieeexplore.ieee.org/abstract/document/1332560.

43. Livak, K. J. & Schmittgen, T. D. Analysis of relative gene expression data using real-time quantitative PCR and the 2(-Delta Delta C(T)) Method. Methods 25, 402–408 (2001).

44. Wang, R. et al. Long nonlZIcoding RNA HOX transcript antisense RNA promotes expression of 14lZI3lZI3σ in nonlZIsmall cell lung cancer. Experimental and Therapeutic Medicine 14, 4503–4508 (2017).

45. Camlin, N. J. et al. Maternal Smoke Exposure Impairs the Long-Term Fertility of Female Offspring in a Murine Model1. Biology of Reproduction 94, 39, 1–12 (2016).

46. Mays, C. E. Effects of Sidestream Smoke on Pregnant Mice and their Offspring. Proceedings of the Indiana Academy of Science 95, 529–536 (1985).

47. Esposito, E. R., Horn, K. H., Greene, R. M. & Pisano, M. M. An animal model of cigarette smoke-induced in utero growth retardation. Toxicology 246, 193–202 (2008).

48. Metzger, M. J., Halperin, A. C., Manhart, L. E. & Hawes, S. E. Association of Maternal Smoking during Pregnancy with Infant Hospitalization and Mortality Due to Infectious Diseases. Pediatr Infect Dis J 32, e1– e7 (2013).

49. Agrawal, A. et al. The Effects of Maternal Smoking During Pregnancy on Offspring Outcomes. Prev Med 50, 13 (2010).

50. Janbazacyabar, H. et al. The Effects of Maternal Smoking on Pregnancy and Offspring: Possible Role for EGF? Frontiers in Cell and Developmental Biology 9, (2021).

51. Suter, M. A. & Aagaard, K. M. The impact of tobacco chemicals and nicotine on placental development. Prenat Diagn 40, 1193–1200 (2020).

52. Suter, M. et al. Maternal tobacco use modestly alters correlated epigenome-wide placental DNA methylation and gene expression. Epigenetics 6, 1284–1294 (2011).

53. Luck, W., Nau, H., Hansen, R. & Steldinger, R. Extent of nicotine and cotinine transfer to the human fetus, placenta and amniotic fluid of smoking mothers. Dev Pharmacol Ther 8, 384–395 (1985).

54. Beyer, D., Mitfessel, H. & Gillissen, A. Maternal smoking promotes chronic obstructive lung disease in the offspring as adults. Eur J Med Res 14, 27–31 (2009).

55. Toppila-Salmi, S. et al. Maternal smoking during pregnancy affects adult onset of asthma in offspring: a follow up from birth to age 46 years. European Respiratory Journal 55, (2020).

56. Gilliland, F. D., Li, Y.-F. & Peters, J. M. Effects of Maternal Smoking during Pregnancy and Environmental Tobacco Smoke on Asthma and Wheezing in Children. Am J Respir Crit Care Med 163, 429–436 (2001).

57. Neuman, Å. et al. Maternal smoking in pregnancy and asthma in preschool children: a pooled analysis of eight birth cohorts. Am J Respir Crit Care Med 186, 1037–1043 (2012).

58. Hayatbakhsh, M. R. et al. Maternal smoking during and after pregnancy and lung function in early adulthood: a prospective study. Thorax 64, 810–814 (2009).

59. Meyer, K. F. et al. The fetal programming effect of prenatal smoking on Igf1r and Igf1 methylation is organ- and sex-specific. Epigenetics 12, 1076–1091 (2017).

60. Lin, P.-I., Shu, H. & Mersha, T. B. Comparing DNA methylation profiles across different tissues associated with the diagnosis of pediatric asthma. Sci Rep 10, 151 (2020).

61. Song, F. et al. Tissue specific differentially methylated regions (TDMR): Changes in DNA methylation during development. Genomics 93, 130–139 (2009).

62. Novakovic, B. et al. Postnatal stability, tissue, and time specific effects of AHRR methylation change in response to maternal smoking in pregnancy. EpigeneticsflJ: official journal of the DNA Methylation Society 9, 377–386 (2014).

63. Armstrong, D. A., Lesseur, C., Conradt, E., Lester, B. M. & Marsit, C. J. Global and gene-specific DNA methylation across multiple tissues in early infancy: implications for children’s health research. FASEB J 28, 2088–2097 (2014).

64. Zhou, J. et al. Tissue-specific DNA methylation is conserved across human, mouse, and rat, and driven by primary sequence conservation. BMC Genomics 18, 724 (2017).

65. Wan, J. et al. Characterization of tissue-specific differential DNA methylation suggests distinct modes of positive and negative gene expression regulation. BMC Genomics 16, 49 (2015).

66. Herzog, E. M. et al. The tissue-specific aspect of genome-wide DNA methylation in newborn and placental tissues: implications for epigenetic epidemiologic studies. Journal of Developmental Origins of Health and Disease 12, 113–123 (2021).

67. Herzog, E. et al. Tissue-specific DNA methylation profiles in newborns. Clinical epigenetics 5, (2013).

68. Chellian, R. et al. Rodent models for nicotine withdrawal. J Psychopharmacol 35, 1169–1187 (2021).

69. McLaughlin, I., Dani, J. A. & De Biasi, M. Nicotine Withdrawal. in The Neuropharmacology of Nicotine Dependence (eds. Balfour, D. J. K. & Munafò, M. R.) 99–123 (Springer International Publishing, 2015). doi:10.1007/978-3-319-13482-6_4.

70. Paolini, M. & De Biasi, M. Mechanistic insights into nicotine withdrawal. Biochemical Pharmacology 82, 996–1007 (2011).

71. Jacobsen, L. K. et al. Effects of smoking and smoking abstinence on cognition in adolescent tobacco smokers. Biological Psychiatry 57, 56–66 (2005).

72. Tager, I. B., Weiss, S. T., Muñoz, A., Rosner, B. & Speizer, F. E. Longitudinal Study of the Effects of Maternal Smoking on Pulmonary Function in Children. New England Journal of Medicine 309, 699–703 (1983).

73. McEvoy, C. T. & Spindel, E. R. Pulmonary Effects of Maternal Smoking on the Fetus and Child: Effects on Lung Development, Respiratory Morbidities, and Life Long Lung Health. Paediatric Respiratory Reviews 21, 27–33 (2017).

74. Goksör, E., Åmark, M., Alm, B., Gustafsson, P. M. & Wennergren, G. The impact of pre- and post-natal smoke exposure on future asthma and bronchial hyper-responsiveness. Acta Paediatrica 96, 1030–1035 (2007).

75. Xiao, R., Noël, A., Perveen, Z. & Penn, A. L. In utero exposure to second-hand smoke activates pro-asthmatic and oncogenic miRNAs in adult asthmatic mice. Environ Mol Mutagen 57, 190–199 (2016).

76. Leon Hsu, H.-H., et al. Prenatal Particulate Air Pollution and Asthma Onset in Urban Children. Identifying Sensitive Windows and Sex Differences. Am J Respir Crit Care Med 192, 1052–1059 (2015).

77. Xiao, R. et al. In utero exposure to second-hand smoke aggravates the response to ovalbumin in adult mice. Am J Respir Cell Mol Biol 49, 1102–1109 (2013).

78. Noël, A. et al. Sex-specific lung functional changes in adult mice exposed only to second-hand smoke in utero. Respir Res 18, 104 (2017).

79. Nielsen, H. C. Androgen receptors influence the production of pulmonary surfactant in the testicular feminization mouse fetus. J Clin Invest 76, 177–181 (1985).

80. Torday, J. S. & Nielsen, H. C. The Sex Difference in Fetal Lung Surfactant Production. Experimental Lung Research 12, 1–19 (1987).

81. Carey, M. A. et al. The impact of sex and sex hormones on lung physiology and disease: lessons from animal studies. American Journal of Physiology-Lung Cellular and Molecular Physiology 293, L272–L278 (2007).

82. Cunningham, J., Dockery, D. W. & Speizer, F. E. Maternal Smoking during Pregnancy as a Predictor of Lung Function in Children. American Journal of Epidemiology 139, 1139–1152 (1994).

83. Drummond, D. et al. Combined Effects of in Utero and Adolescent Tobacco Smoke Exposure on Lung Function in C57Bl/6J Mice. Environ Health Perspect 125, 392–399 (2017).

84. Singh, S. P. et al. Maternal Exposure to Secondhand Cigarette Smoke Primes the Lung for Induction of Phosphodiesterase-4D5 Isozyme and Exacerbated Th2 Responses: Rolipram Attenuates the Airway Hyperreactivity and Muscarinic Receptor Expression but Not Lung Inflammation and Atopy. The Journal of Immunology 183, 2115–2121 (2009).

85. Maideen, N. M. P. Tobacco smoking and its drug interactions with comedications involving CYP and UGT enzymes and nicotine. World Journal of Pharmacology 8, 14–25 (2019).

86. Anttila, S. et al. Methylation of Cytochrome P4501A1 Promoter in the Lung Is Associated with Tobacco Smoking. Cancer Research 63, 8623–8628 (2003).

87. Hankinson, O. The aryl hydrocarbon receptor complex. Annu Rev Pharmacol Toxicol 35, 307–340 (1995).

88. Kim, J. Expression of cytochromes P450 1A1 and 1B1 in human lung from smokers, non-smokers, and ex-smokers. Toxicology and Applied Pharmacology 199, 210–219 (2004).

89. Rendic, S. & Di Carlo, F. J. Human cytochrome P450 enzymes: a status report summarizing their reactions, substrates, inducers, and inhibitors. Drug Metab Rev 29, 413–580 (1997).

90. Okey, A. B. Enzyme induction in the cytochrome P-450 system. Pharmacology & Therapeutics 45, 241–298 (1990).

91. Ohashi, H. et al. The aryl hydrocarbon receptor–cytochrome P450 1A1 pathway controls lipid accumulation and enhances the permissiveness for hepatitis C virus assembly. J Biol Chem 293, 19559– 19571 (2018).

92. Rendic, S. & Guengerich, F. P. Contributions of Human Enzymes in Carcinogen Metabolism. ACS Publications https://pubs.acs.org/doi/pdf/10.1021/tx300132k (2012) doi:10.1021/tx300132k.

93. Nebert, D. W. et al. Role of the aromatic hydrocarbon receptor and [Ah] gene battery in the oxidative stress response, cell cycle control, and apoptosis. Biochemical Pharmacology 59, 65–85 (2000).

94. Uno, S., Sakurai, K., Nebert, D. W. & Makishima, M. Protective role of cytochrome P450 1A1 (CYP1A1) against benzo[a]pyrene-induced toxicity in mouse aorta. Toxicology 316, 34–42 (2014).

95. Uno, S. et al. Oral Benzo[a]pyrene in Cyp1 Knockout Mouse Lines: CYP1A1 Important in Detoxication, CYP1B1 Metabolism Required for Immune Damage Independent of Total-Body Burden and Clearance Rate. Mol Pharmacol 69, 1103–1114 (2006).

96. Neuhold, L. A., Shirayoshi, Y., Ozato, K., Jones, J. E. & Nebert, D. W. Regulation of mouse CYP1A1 gene expression by dioxin: requirement of two cis-acting elements during induction. Molecular and Cellular Biology 9, 2378–2386 (1989).

97. Gluckman, P. D., Hanson, M. A. & Low, F. M. Evolutionary and developmental mismatches are consequences of adaptive developmental plasticity in humans and have implications for later disease risk. Philosophical Transactions of the Royal Society B: Biological Sciences 374, 20180109 (2019).

98. Frankenhuis, W. E., Nettle, D. & McNamara, J. M. Echoes of Early Life: Recent Insights From Mathematical Modeling. Child Development 89, 1504–1518 (2018).

99. Godfrey, K. M., Lillycrop, K. A., Burdge, G. C., Gluckman, P. D. & Hanson, M. A. Epigenetic Mechanisms and the Mismatch Concept of the Developmental Origins of Health and Disease. Pediatr Res 61, 5–10 (2007).

100. Marwick, J. A. et al. Cigarette Smoke Alters Chromatin Remodeling and Induces Proinflammatory Genes in Rat Lungs. Am J Respir Cell Mol Biol 31, 633–642 (2004).

101. Meldrum, D. R. et al. Adaptive and maladaptive mechanisms of cellular priming. Ann Surg 226, 587– 598 (1997).

102. Kentner, A. C., Cryan, J. F. & Brummelte, S. Resilience Priming: Translational models for understanding resiliency and adaptation to early-life adversity. Dev Psychobiol 61, 350–375 (2019).

103. Hales, C. N. & Barker, D. J. P. The thrifty phenotype hypothesis: Type 2 diabetes. British Medical Bulletin 60, 5–20 (2001).

104. Gluckman, P. D. & Hanson, M. A. Living with the Past: Evolution, Development, and Patterns of Disease. Science 305, 1733–1736 (2004).

105. Hsu, C.-N. & Tain, Y.-L. Animal Models for DOHaD Research: Focus on Hypertension of Developmental Origins. Biomedicines 9, 623 (2021).

106. Lavigne, É. et al. Maternal exposure to ambient air pollution and risk of early childhood cancers: A population-based study in Ontario, Canada. Environ Int 100, 139–147 (2017).

107. Erickson, A. C. & Sbihi, H. Biological embedding, the air we breathe, and carcinogenesis. The Lancet Planetary Health 2, e149–e150 (2018).

108. Tomczyk, M. M. et al. Mitochondrial Sirtuin-3 (SIRT3) Prevents Doxorubicin-Induced Dilated Cardiomyopathy by Modulating Protein Acetylation and Oxidative Stress. Circulation: Heart Failure 15, e008547 (2022).

